# Using iCn3D and the World Wide Web for structure-based collaborative research: Analyzing molecular interactions at the root of COVID-19

**DOI:** 10.1101/2020.07.01.182964

**Authors:** Philippe Youkharibache, Raul Cachau, Tom Madej, Jiyao Wang

## Abstract

The COVID-19 pandemic took us ill-prepared and tackling the many challenges it poses in a timely manner requires world-wide collaboration. Our ability to study the SARS-COV-2 virus and its interactions with its human host in molecular terms efficiently and collaboratively becomes indispensable and mission-critical in the race to develop vaccines, drugs, and neutralizing antibodies. There is already a significant corpus of 3D structures related to SARS and MERS coronaviruses, and the rapid generation of new structures demands the use of efficient tools to expedite the sharing of structural analyses and molecular designs and convey them in their native 3D context in sync with sequence data and annotations. We developed iCn3D (pronounced “I see in 3D”) ^1^ to take full advantage of web technologies and allow scientists of different backgrounds to perform and share sequence-structure analyses over the Internet and engage in collaborations through a simple mechanism of exchanging “lifelong” web links (URLs). This approach solves the very old problem of “sharing of molecular scenes” in a reliable and convenient manner. iCn3D links are sharable over the Internet and make data and entire analyses findable, accessible, and reproducible, with various levels of interoperability. Links and underlying data are FAIR ^2^ and can be embedded in preprints and papers, bringing a 3D live and interactive dimension to a world of text and static images used in current publications, eliminating at the same time the need for arcane supplemental materials. This paper exemplifies iCn3D capabilities in visualization, analysis, and sharing of COVID-19 related structures, sequence variability, and molecular interactions.

## INTRODUCTION

With the COVID-19 pandemic our ability to study the virus and virus-host interactions in-depth and collaboratively has become extremely important. We already know key SARS-COV-2 viral proteins at the molecular level and some of the molecular interactions that allow the virus spike to bind its human host ACE2 receptor. Structural analyses have become de facto mission-critical for the development of new (or repurposed) drugs, vaccines, or antibodies, and making them instantaneously available worldwide is imperative. For that to occur we need to lower the barrier of entry to study molecular structures for scientists that are not trained in that field and enable the discovery process and sharing of analyses in a self-teaching environment.

Structure-based antigen design, computational biology, and protein engineering provide methods to make vaccines with speed and precision ^3^. This is a reason for hope in developing a vaccine in a short time frame. Structure-based drug design, whether on small molecules or monoclonal antibodies, provides a pathway to possible treatments. The global need for vaccines and drugs and the wide geographic diversity of the pandemic require more than one effective vaccine or drug design approach, and the full development pathway for an effective vaccine for SARS-CoV-2 requires the collaboration of industry, government, and academia at an unprecedented scale ^4^. “Stopping the pandemic could rely on breakneck efforts to visualize SARS-CoV-2 proteins and use them to design drugs and vaccines” ^5^. Yet, **the current tools are limited in their ability to exchange information at the required level.** iCn3D offers an initial contribution in that direction, by making the sharing and collaboration on structure and structure analysis possible, peer to peer, and through preprint and publication channels, seamlessly.

With the COVID-19 pandemic, an avalanche of new experimental and modeled structures became available in a very short time over the web, and the production of new structures is accelerating. In one month the number of structures has almost doubled (https://www.ncbi.nlm.nih.gov/structure?term=SARS-COV-2). It took only a few weeks after the publication of the SARS-COV-2 virus genome sequence to get the first 3D structures of the virus spike interacting with the human ACE2 receptor (see <u>gallery</u>), and new experimental 3D structures are produced at an unprecedented rate. Many are available as 3D coordinates in repositories and their descriptions/annotations are spread in a myriad of papers or preprints on the Internet. These structures are the basis of a very large number of structural analyses, modeling efforts, and structure-based design projects all over the planet: on vaccines, on broadly neutralizing antibodies, and on drug lead explorations.

Yet, structural information is still exchanged in the year 2020, for the most part, as it was decades years ago. Structures are still shared as arcane sets of 3D coordinates and are interpreted by sophisticated, often proprietary software applications, some outdated or not properly maintained. To this date, structural annotations are still lengthy textual descriptions and represented as 2D pictures in papers, and sometimes through supplemental videos. While structural biologists and molecular modelers, armed with extensive knowledge in molecular structure and long experience in using complex proprietary software to process data, may be comfortable with the status quo, structural analyses remain opaque to a broad range of scientists. To make matters worse, and especially in the case of COVID-19, structural annotations are dispersed in hundreds of papers, supplements, preprints, and now archives.

By contrast, it only takes a second to share a structural annotation worldwide through 3D web links ^1^, exactly as one would share a Google map or a Google Doc. For example, a simple link can focus on <u>the SARS-COV-2 Receptor Binding domain interacting with the ACE2 receptor</u> (Link 1), or allow one to visualize and analyze epitopes used by neutralizing <u>antibodies against SARS-COV-1 vs. SARS-COV-2</u> (Link 2), or analyze where <u>mutations may occur across coronaviruses</u> (Link 3) ***in a structural context***.

The unprecedented volume of new structures and new sequencing data shared through the Internet as data sets, preprints and publications exacerbates the need to perform structural analyses quickly and share these results with collaborators and through online publications. The role of 3D in this fast moving field is critical because it is the one form of annotation that can effectively organize and coordinate the others. In other words, we can correct a faulty 1D alignment or improve the interpretation of a sequence pattern if the 3D (structure) information can be adequately harnessed. iCn3D has been designed for that purpose ***and*** it can help foster collaborative research over the World Wide Web. It does not replace any modeling or expert tool. It emerges from a new paradigm that emphasizes visualization, analysis, and sharing of sequence-structure information over the web.

## RESULTS AND DISCUSSION

The simplest way to look at iCn3D, beyond simply visualizing 3D structures of molecular structures, is to see it as a tool to investigate the structure of molecular assemblies ***interactively and cooperatively***. iCn3D, literally, allows the interaction with molecular objects: highlighting, delineating and extracting elementary structural properties at any level of detail (atom, residue, domain, chain, or any arbitrary set of residues or atoms) to reveal the structural underpinning of a given molecular structure and constituent molecular interactions. These are all very simple and very well known descriptive elements (in a classical context): covalent bonds, Hydrogen-bonds, non-bonded interactions (Van der Waals). The strength of the program is its ability to analytically address any set of such elements in isolation, collectively and in interaction.

In order to convey the power of interactive structural analysis, we investigate a molecular system of current interest: the SARS-COV spike proteins, and their interactions with the human ACE2 receptor in their initial (pre-fusion) binding stage, and highlight the analytical methods employed.

This exploration is reported as an investigation, not a tutorial, yet at every step the reader will be offered a 3D link that encapsulates a reproducible analysis within the context of the analytical software itself, to make the experience a ***self-teaching*** one. Through these links, the reader will be offered a graphical description of the analysis performed, the tool to reproduce that analysis, and very importantly the ability to go beyond in extending a proposed analysis, making new observations, new connections, new inferences. The power of these links is in sharing and disseminating analyses with peers to open collaborative research. Currently, these links have been used mostly in education, where professors can teach their students about molecules and molecular interactions, concentrating on chemistry or physics of interactions, not on how to use the software. This is mostly a descriptive (passive) use. The link is live and an invitation to active, interactive use of molecular objects. Exchanging links between scientists is in essence a mechanism for collaborative, iterative, and discursive analysis.

These URLs are “lifelong” and can be accessed from a preprint or a publication. We started to publish 3D analyses as links ^1,6^. This paper further demonstrates its use with links that annotate structures and molecular interactions between SARS-COV-2 and the human ACE2 receptor, or with antibodies. Each iCn3D link can be reused, modified, extended, saved, and reshared, making a paper not an end but potentially a new beginning.

All SARS-COV-2 structures used in this study were deposited by their authors in the PDB database. They are accessible as initial iCn3D links on the <u>NCBI Structure website</u>. In addition, unpublished structures can be loaded in a PDB or MMCIF format ^7^, and exchanged as an “iCn3D PNG image” file including data and annotations; they can also be shared through a web server privately on an intranet or publicly through the Internet. We have just released iCn3D version 2.17.5 to help analyze, annotate, and share sequences & structures. The software is open-source and available from <u>GitHub</u> as an invitation to software developers to join in.

In the following sections, we will exemplify iCn3D’s 3 main functions:

1. Molecular Visualization.
2. Interactive Structural Analysis.
3. Sharing, Collaborating, Publishing.

As mentioned before, this will be reported as an investigation on SARS-CoV/ACE2 interaction analysis, not a software description or a tutorial, in order to make the experience a ***self-teaching*** one.

### A brief summary of the analysis developed in the Methods section

We start by observing the architecture of the spike trimer vs. the ACE2 receptor, focusing on the Receptor Binding Domain (RBD)-ACE2 interaction and get deeper and deeper in the analysis of the details of molecular interactions through a set of annotated links that can be manipulated by the reader.

Simultaneous visualization of the Spike chain sequences (1D), domain organization, and 3D structure allow zooming on regions of interest. We can observe a spike trimer exhibiting different conformations of its three RBDs. Beyond, since experimental structures usually focus on a limited number of domains, chains, or assemblies, one can use them to build larger composite assemblies through partial overlaps (domain level structural alignment). In doing so, we can visualize for example an ACE2 [quasi] intact dimer with one or two spikes binding one or two of the ACE2 peptidase domains.

We then drill down, looking at the binding interface between SARS-COV-2 Spike’s RBD and the ACE2 receptor in terms of residue interactions. We compare CoV-1 vs. CoV-2 RBDs in binding ACE2 (differential binding analysis). We then look at sequence variants across beta Coronaviruses RBDs, including MERS, analyzing conservation in the Spike RBD, comparing animal and human sequences. Since iCn3D accesses seamlessly dbSNP, we map human ACE2 SNPs to see how polymorphism may act on structure, especially at the viral interface. Neutralizing antibodies are an important focus in COVID-19 research and some structures are available, we look at some that bind RBD epitopes and how these compare with the ACE2 binding RBD surfaces. Glycans on the virus surface or within the ACE2 receptor are naturally highly studied and will be observed.

### 3D links are used along the way

Not only do links present the results of analyses, and share the data and annotations with readers, they offer a starting point for challenging analyses and performing a further investigation. This is the 3rd function of iCn3D: providing a capability to share structural analysis and annotations with collaborators and peers and use them in publications. These links are in fact already used today by a number of professors in teaching about protein structure and molecular interactions, and we envision them as being used for peer review and publication, as we have started to do ourselves in our latest preprints and papers ^1,6^.

## METHODS

### 1. Molecular Visualization

An initial goal in developing iCn3D was to visualize biological molecules’ structures to help unveil the structural basis of biological function. In molecular terms, it meant the ability to study interacting biological molecules with the appropriate molecular representations. To help guide this development, we defined “Twelve Elements of visualization and analysis for the tertiary and quaternary structure of biological molecules” ^8^, that are now well underway in their implementation. We will review here the most relevant elements.

#### 3D data retrieval

Currently, the retrieval of protein 3D coordinates is essentially the same as it has ever been since the inception of the protein data bank (www.pdb.org) in Brookhaven National Labs, almost fifty years ago ^9^. The use of a “PDB id” and the maintenance of the PDB data repository since 1971 has been a wonderful means of data sharing. This is also the starting point for any structural analysis with any public or commercial software, such as Chimera, etc. 3D coordinates stored in a PDB file (whether old or newer style) are the basic data. With the introduction of iCn3D, one can start directly from a link that gives access to a 3D visual representation associated with a 1D sequence window and a 2D schematic representation. These are available for any PDB structure from the NCBI Structure web site (www.ncbi.nlm.nih.gov/structure) with a good number of annotations that are generated automatically such as conserved protein domains, intermolecular interactions, dbSNP or ClinVar annotations, secondary structures, or functional sites.

One can also visualize any structure from a user-defined coordinate file in a PDB (or MMCIF) format. Users can create their own repository for use on an intranet or on the Internet (see an example at the Frederick National Lab who is serving <u>verified COVID-19 related structures</u> correcting issues that may be found in recently deposited SARS-COV-2 structures).

#### “Set Selection” and synchronized 1D/2D/3D visualization

iCn3D has the ability to name any molecular entity at any level of detail (chain, domain, supersecondary structure, residue, atom). This gives great flexibility for analysis, especially in studying molecular interactions. Practically, the analytical power of iCn3D relies on its ability to select sets of molecular entities at any level of detail and to name them. It allows us to study and compare sets across homologous (or non-homologous) families and very important to study their interactions, as will become clear in the following. Sets open the door to virtually unlimited flexibility since they can be combined through Boolean operations. That flexibility and breadth of analysis will grow even more as we introduce more and more molecular properties, while each property alone or in combination can enable set selection (This is a key design Element #3 - ^8^)

We have implemented a parallel, synchronized visualization of 1D sequence and 3D structure visualization. This is of tremendous use to follow sequences in their (3D-folded) operational environment. We have added 2D, which allows simple schematic representations and projections. This is, surprisingly, an untapped realm of visualization, and we are just scratching the surface at the moment. 2D is particularly useful to simplify and illustrate molecular interactions. 2D illustrations are well suited for paper publications. Meanwhile, the 3D is only perceivable through 3D visualization, and snapshots of a 3D screen used in papers to this date usually do not convey the depth of information they are meant to represent. To achieve this we use 3D links, which by the same token allow simultaneous use and visualization in 1D 2D 3D. In addition, data can be represented as Tables that list residues/atoms properties, such as solvent accessibility, contacts/pairwise interactions. The latter can be projected on 3D structures, or exported for external analysis. Conversely, externally computed data can be imported for visualization and analysis within iCn3D.

#### i Cn3D triple focus: Visualization, Structural Analysis, Sharing

It is important to understand what iCn3D does and does not do. It was designed to enable visualization and analysis of structures and molecular interactions at any level of detail in a straightforward manner, as we shall illustrate in the following. Very importantly it is designed to share molecular scenes and entire structural analyses in 1D, 2D, 3D, and Tables (containing numerical data) to enable interactive collaborative research and data sharing. The simple use of links allows both passive visualization and/or active analysis. These links can be shared through emails, papers, preprints, courses, etc. i.e. in any live document. The software is centered on structural visualization, analysis, and sharing. It does not perform modeling or complex computational biology simulations. It can however import and analyze structures determined experimentally or computationally, and molecular properties that may be produced through computational biology.

#### Aword on Reference numbering

This section can be skipped until the reader comes across with difficulties in dealing with residue numbers. **Residue numbering** may seem a mundane topic, yet it is a major unresolved issue that could be resolved in any given context, and that would save countless hours of tedious work (and tedious reading!) to scientists analyzing sequences and structures. In fact, this problem could be discussed at length, and an entire symposium may not suffice to cover all the issues. **There is no reference numbering used on any protein**, except to some extent on antibodies with the simple and brilliant idea of Kabat ^10^ to number residues in order to compare positions, presented over 30 years ago. Naturally, with hypervariable regions and insertions, numbers may need extensions such as 100A, B, etc., as it happened in antibody CDR3 loop regions, but that numbering has endured, and some international organizations maintain databases with a reference numbering on antibodies ^11^. Using a reference numbering can be tricky in widely variable systems before these systems are known well enough of course, but a reference is better than none. The field of genomics has this incredible advantage of using Reference Genomes with a genome reference numbering, at least on an organism basis. One can think of what would be the impossible task of genome assembly without a reference numbering to relate to!

In dealing with proteins, the community should adopt a reference numbering at least for widely studied problems such as SARS Coronaviruses proteins and receptors, certainly spikes and certainly for RBDs as we will need to compare many species receptors and many viral strains. Already too many preprints on SARS-CoV-1 and SARS-CoV-2 are using different numbers for the same residue, for example as we shall see Y453 <> Y440, and we add some more numbering gymnastics in using relative numbers for each Y135<>Y114. This is a problem we can solve on our end, yet ultimately it is a community issue to adopt a single reference numbering. For the ACE2 receptors, we are lucky as ACE2 is a highly conserved protein with an (almost) constant number of 805 residues across all species, yet in the literature, as for any protein, numbers differ when one considers the signal peptide or not, and even when one considers the starting Methionine or not in numbering. The solution would be to use numbers according to REFSEQ ^12^ or UNIPROT ^13^ protein sequences. Structural Biologists have often used numbers that match these, but of course, many structures in the PDB predate all these sequence databases.

iCn3D’s primary source is MMDB ^14^ that uses an internal numbering starting at 1 for the first residue read in and then numbered consecutively. For example in iCn3D ACE2 numbers will start at position 1, which is position 19 in reference sequence in UNIPROT <u>Q9BYF1</u>, same transcript as REFSEQ <u>NP_068576.1</u>. In the general case, things may be more complex. For example, REFSEQ has two transcripts one shorter one longer associated with the <u>ACE2 gene</u>. For a general reference numbering solution in proteins, the problem can become quite complex with alternative splicing and multiple transcripts associated with a gene. Historically the problem of numbering in sequence analysis has not been seen as a forefront issue since sequence mappings have been used, not numbers. Yet when analyzing sequences and structures and interactions that can change in sequence but maintain geometric relationships, a number base is unavoidable, and it can be made much simpler and reliable with a reference numbering across all protein sequence occurrences from many organisms.

iCn3D keeps track of the original PDB numbering that is available in the “sequence and annotations window”. If the data is loaded directly from the native PDB structure file, then native PDB numbers are used. All these numbering issues are historical. Anyone who has used PDB structures knows far too many issues in numbering, with missing residues, insertions, deletions, etc. Yet going forward, for important collaborative projects such as those rapidly emerging due to the fast paced efforts triggered by the COVID-19 pandemic, the community should adopt a reference sequence numbering to be used across all species and all strains. We do not solve the problem in this version of the program but we will have to solve it. So our apologies if we use multiple numbers to refer to a single position across structures of SARS-CoV-1, SARS-CoV-2, and MERS … from different strains or for infected species. This is not a limitation of iCn3D but what happens and will happen in virtually any published paper until a reference numbering system is adopted.

### 2. Structural Analysis

#### Visualization of the SARS-COV-2 protein spike

##### Protein domains delineation, visualization, and coloring

The Virus Spike structure has been determined by Electron Microscopy (EM) (^15^,^16^). A first inspection consists of delineating and visualizing the various domains that compose the spike. In Figure 1.A, we can visualize the spike trimer where one monomer is shown as ribbons with individual domains colored, in particular, the Receptor Binding domain in Cyan, and the N-terminal domain (NTD) in dark blue. Since we need to study the spike interaction with its receptor, we can bring this spike in its larger structural context, combining various structures determined experimentally, look at the spike trimer in context.

**Figure 1.**
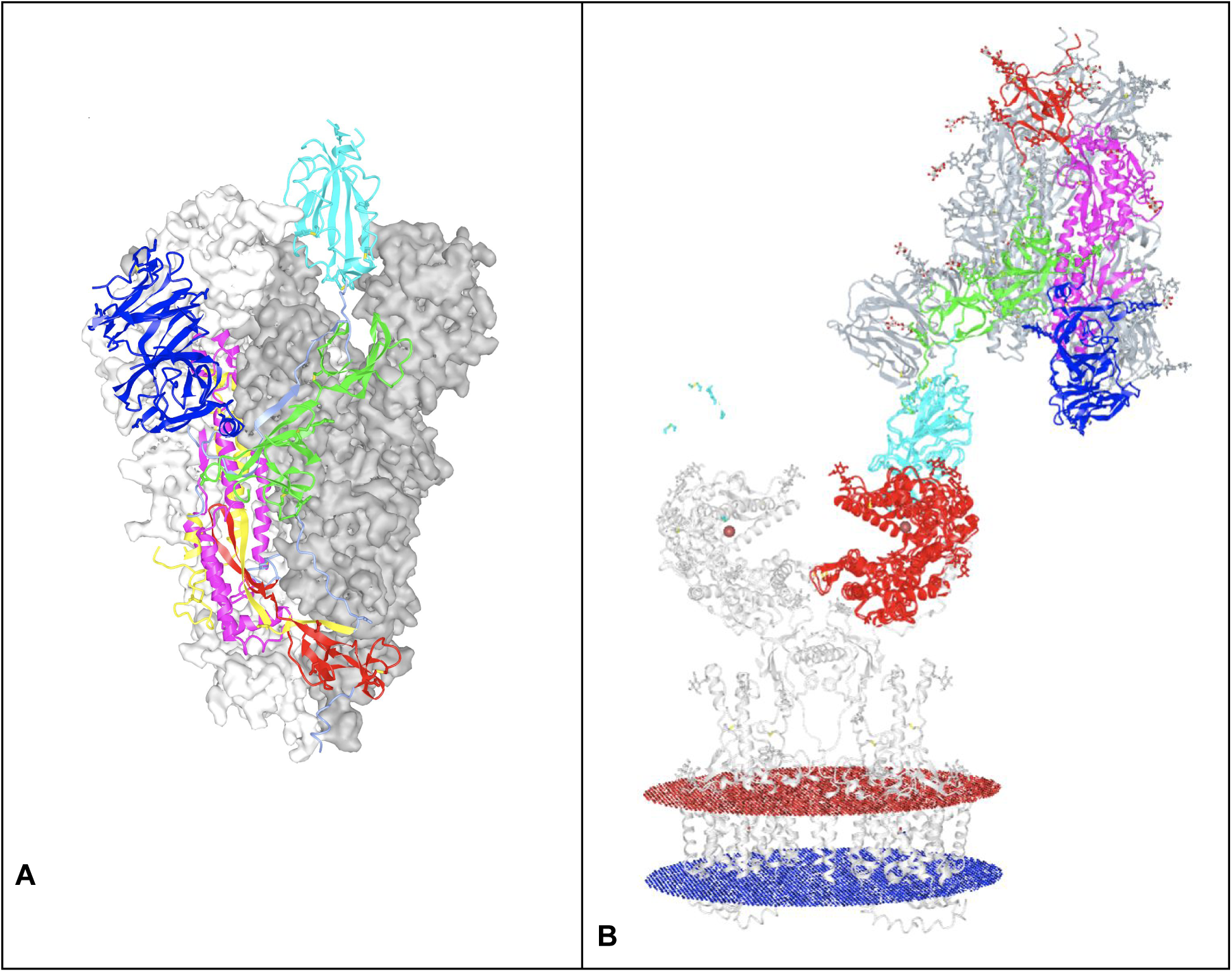
SARS-COV spike. **A) SARS-COV-2 Trimer with one monomer showing <u>domain composition</u>** <u>in ribbon representation</u> (Link 4) (PDBid 6VSB) with domains colored for one monomer: the NTD in dark blue and the RBD in Cyan. The other two monomers are represented as molecular surfaces in white and grey. <u>SARS-COV-1</u> <u>spike is highly homologous and can be superimposed for the whole trimer (PDB 6CS2)</u> (Link 5), less the mobile RBD depending on the up/down conformations **B) Building a Structural assembly through matching domain alignment within 2 PDB structures: Visualizing a whole spike binding to ACE2.** Shown here is <u>the spike Trimer structure (6CS2) of</u> **<u>SARS-COV-1</u>** <u>binding the full-length ACE2 peptidase (PDB 6M17)</u> (Link 6) aligned with the ACE2 peptidase domain. Notice the match, just on that domain alignment between RBDs of SARS-CoV-2 (6M17) and SARS-COV-1. The overall RBD-ACE2 interface is a remarkably conserved SARS-CoV-2 vs. SARS-CoV-1. The intact ACE2 receptor dimer is complexed with B0AT1 transporter. The cell membrane is indicated in red and blue planar boundaries. The RBDs are colored Cyan as in A) and in dark blue for the monomer chain, the other chains of the spike trimer two in gray

##### Building and visualization Biological Assemblies: Spike(s) vs. intact ACE2 dimer?

Structural Biology (XRAY, NMR, EM …) provides structures of domains or chains in isolation and in interaction. Rarely does it offer a full complex, and one may need to combine/assemble a full complex for further analysis. Such an assembly process from parts is commonly used in assembling EM structures while fitting an electron density map. Similarly, when studying a problem one may need to bring together various protein domains in interaction, while these may be spread in multiple “structure files”, each described independently in a paper, and aggregate the parts to get a broader picture.

The SARS-COV-2 spike binding to the human ACE2 receptor through its RBD provides a viral entry point for the COVID-19 infection in humans. A structure of the (quasi) intact ACE2 dimer with the RBD binding has been solved independently ^17^ and we can recompose the assembly for the ACE2-spike trimer binding to the intact ACE2. The composite assembly on one spike vs. one ACE2 monomer receptor is assembled by superposing the RBD that is common to both experimental structures. This is made easy through the use of the VAST+ database, that aligns tertiary and quaternary structures from the PDB ^14,18^, accessible through iCn3D. In Figure 1.B, we can visualize one spike interacting through one RBD (in Cyan) in the up position with one ACE2 peptidase domain (in Red). What transpires from this buildup is that, sterically, an ACE dimer can accommodate the binding by two spikes, one can build a dimeric assembly by simply repeating that same buildup on the second RBD binding the second ACE2 peptidase domain (we did hide in Figure 1B that second RBD which is present in the original structure).

##### Visualizing the SARS-COV-2 RBD binding to its human ACE2 receptor

One can zoom in from the assembly level down to residue and atom level to <u>highlight the interacting residues in sequence (1D) and in 3D</u> (Link 1). Most interactions are situated for all but one in a subdomain of the Receptor Binding Domain (RBD), called the RBM (Receptor Binding Motif) on the virus side vs. binding to the Receptor Binding site (RBS) spread on two subdomains of the ACE2 peptidase domain, most mostly concentrated on the N-term helical subdomain (see Figure 2). In the following we focus on the (RBD) and RBM binding to the human ACE2 receptor ^19 20^. Before we do so, let’s review the architectural features of the RBD itself, common to known beta coronaviruses.

**Figure 2.**
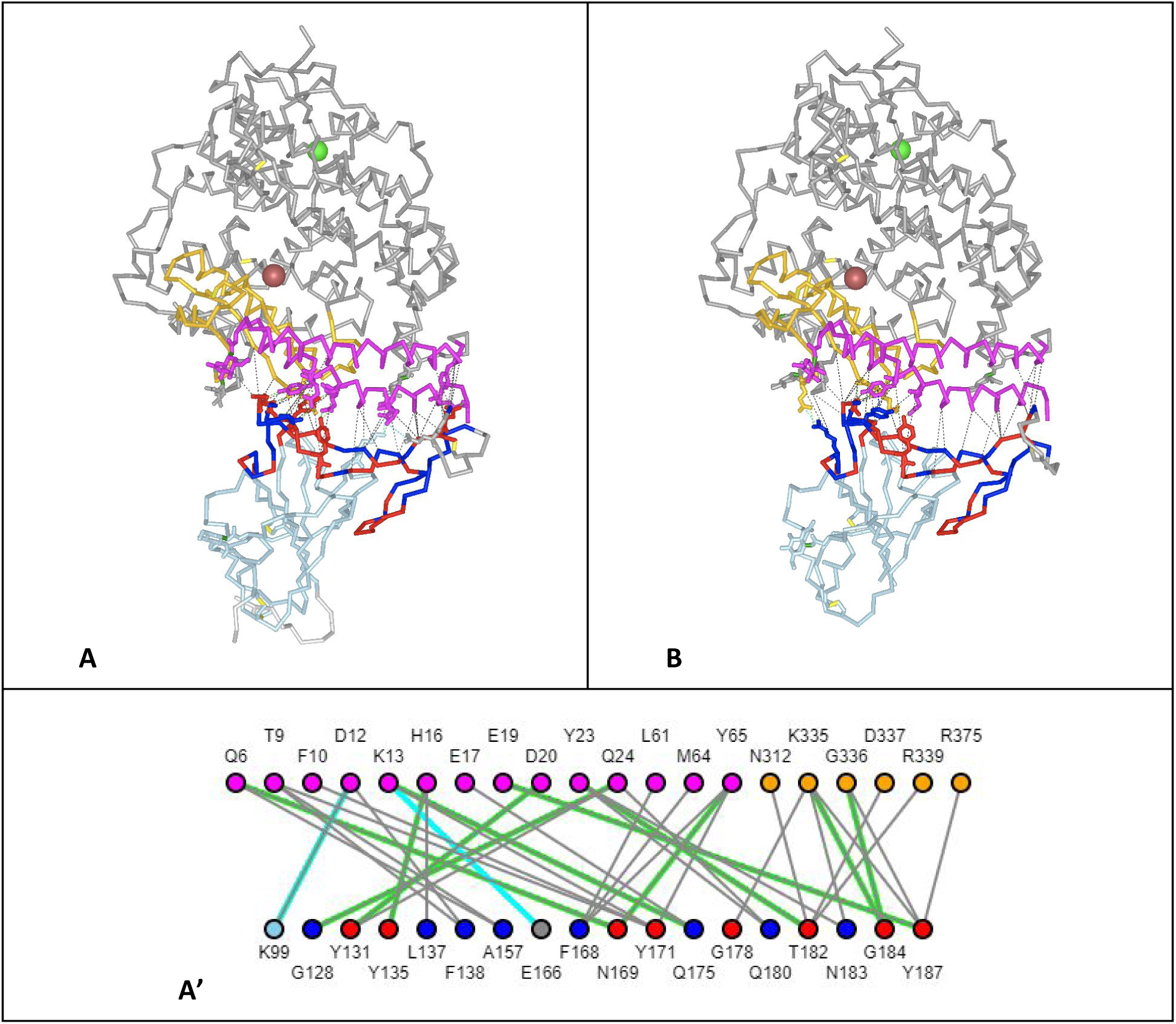
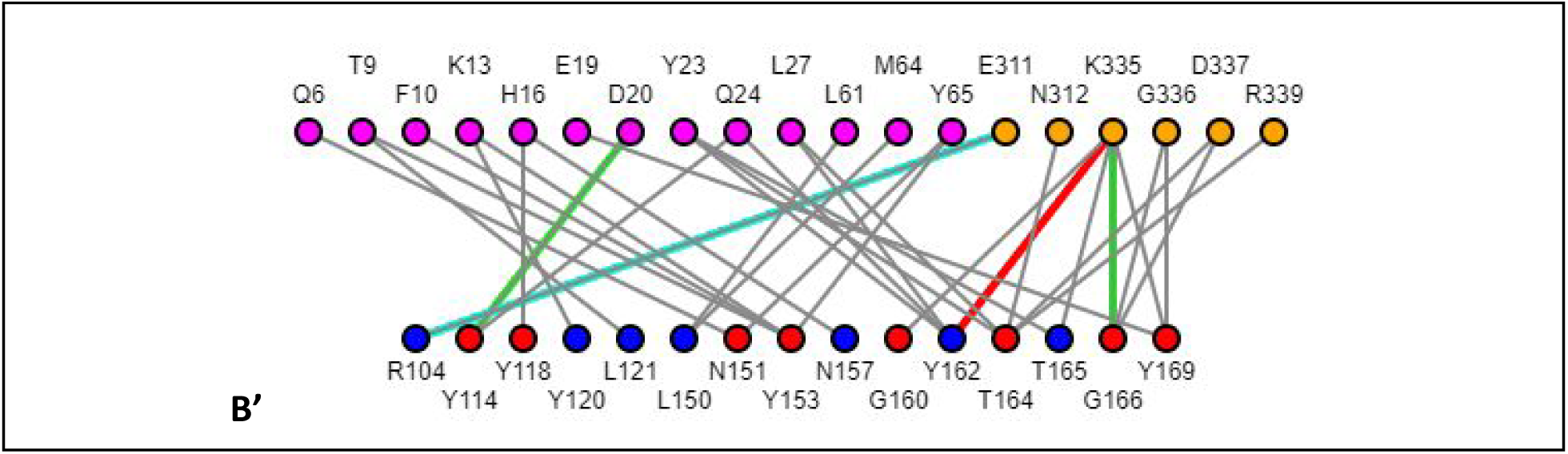
Comparing interacting residue networks at the ACD2-RBD interface (Link 10) **A-A’) for SARS-CoV-2 B-B’) SARS-CoV-1.** ACE2 in grey with ACE2 RBS (Recep[tor Binding Site) is situated in two subdomains, colored magenta for N-terminal helical domain of the ACE2 receptor res#1-65, i.e. #19-83 in ACE2 numbering, the 18-residue difference in numbers come from the removal of the signaling peptide sequence) and orange for its second binding subdomain (res 307-339). Any residue or pair can be selected in the 3D link for detailed visualization and analysis. The nodes of the network are interacting in ACE2 residues vs. the virus RBD/RBM, colored blue, and red when conserved. Interactions of different types are represented by lines between notes (residues) H-bonds (green), Salt Bridges(cyan) between charged residues, Cation-Pi (red), and pi-pi(blue) interactions between aromatic rings. They are computed with various thresholds that can be modified.

#### The architecture of the coronaviruses’ RBD: the RBD framework and the RBM plugin

There are four types of common Coronaviruses that infect humans: 229E (alpha coronavirus), NL63 (alpha coronavirus), OC43 (beta coronavirus), HKU1 (beta coronavirus), and three new beta coronaviruses that appeared recently, originating from animals: MERS-COV, SARS-CoV-1, and SARS-CoV-2, the latter being the origin of the COVID-19 pandemic (https://www.cdc.gov/coronavirus/types.html).

Beta coronaviruses RBDs share the same beta fold framework and vary the RBM subdomain to bind a variety of receptors. Currently, known receptors are peptidases: SARS-CoV-1 and SARS-CoV-2 also bind ACE2. MERS-CoV binds DPP4, with a slightly different RBM. HCoV-229E an alpha coronavirus with yet another, larger RBM binds aminopeptidase N, while receptors for HCoV-OC43 and HKUI, two beta coronaviruses are still unknown. Alpha coronaviruses use a totally different, simpler, RBD with a beta-barrel fold. Intriguingly, NL63 different beta also binds ACE2. (https://www.ncbi.nlm.nih.gov/Structure/cdd/cddsrv.cgi?uid=394827)

If we look at the RBDs of alpha and beta coronaviruses for which we have structures (see Table S1), they all share a common architecture composed of an all beta-sheet **framework** that supports what we would describe as a **beta plugin** RBM (Receptor Binding motif), where the N and C termini of the RBM plug into a beta-hairpin of the framework that would otherwise be a turn/loop. Many protein folds have robust frameworks that accommodate entire domains as plugins. This is in fact a common modular architecture, that is not stressed enough. In principle, any domain with N/C termini joining in close proximity can be plugged-in another domain framework if its folding is robust enough. Incidentally and interestingly, a similar overall RBD architecture is observed for example in other viruses such as the Murine or Feline Leukemia Viruses (pfam00429 TLV_coat ENV polyprotein), albeit with totally different framework or plugin folds. Such a plugin architecture seems to be a brilliant evolutionary technique for viruses to evolve binding motifs that can adapt to different cell surface receptors while maintaining a framework that conserves a needed conformational articulation of the RBD vs. the other spike domains. From a structural alignment of alpha and beta coronaviruses, we seek to determine which residues may be conserved in both sequences and in structure in the RBDs (see Figure 1).

#### Comparing SARS-CoV-1 vs. CoV-2 RBDs in binding ACE2 in iCn3D

##### A simple interface analysis of a complex network of interactions

A protein-protein interface is a very complex web of electronic interactions between residues. XRay structure coordinates provide average positions of atoms that we idealize as points in Euclidean space. This gives us a rather simple representation of points (atoms or residues) and lines (bonds and nonbonded interactions) and we can use Euclidean distances between residues (Carbon-alphas) and/or specific atoms in residues at the interface to describe the interactions between two proteins, in our case between the Spike RBD and its human ACE2 receptor. Most interface descriptions use a cutoff distance of 4A to identify “key interactions”. iCn3D can identify a number of interaction types with variable cutoffs. In the example below (Figure 2), we compare a structure of SARS-CoV-2 (PDB 6M0J) with SARS-CoV-1 (PDB 2AJF) and delineate the corresponding networks of interactions. It is amazing how mutations between the two viruses rewire the interface, yet <u>maintain both the tertiary structure and the overall complex quaternary structure within 0.70A</u> (considering CA only in the alignment over 758 residues with 94% sequence identity overall). This quaternary interface is maintained even with the evolution of residues in the RBD vs. the constant human ACE2 sequence, showing that many variants can bind the same ACE2 RBS. It shows that a dynamic adaptation, rewiring of interactions to the ACE2 sequence-structure. The devil is in the details and one can compare the rewiring of interactions that show an exquisite variety of local variations.

One can characterize and compare SARS-CoV-1 and SARS-CoV-2 RBD interfaces by superimposing the two complexes (quaternary structures) <u>6M0J vs. 2AJF</u> (Link 7) or alternatively <u>6M0J vs. 3SCI</u> (Link 8), and visualize the differences in both sequence and structure simultaneously (see Figure 2). A Superimposed view or a <u>side by side comparison of SARS-CoV-1 and SARS-CoV-2 RBDs</u> (Link 9) shows remarkable overall conservation of the virus-cell receptor complex.

##### Comparing networks of interacting residues in SARS-CoV-1 and SARS-CoV-2 *vs.* ACE2

We can analyze in parallel the two <u>SARS-CoV-1 - ACE2 vs. the SARS-Cov-2 - ACE2</u> (Link 10) interfaces and delineate the interactions, conserved in position and sequence, or position despite sequence differences that create subtle changes in “wiring” difficult to perceive without a detailed analysis, especially some new interactions in SARS-CoV-2 vs. SARS-CoV-1. Figure 2 shows an interaction network representation for each. The 3D links give access to all details of the interactions simultaneously in 3D, 2D, 1D, and Tables, allowing zooming in any residue pair interaction, categorized as H-bonds, Salt Bridges between charged residues, Cation-Pi, and Pi-Pi interactions between aromatic rings. Since XRay or EM structures present irregularities, it is always prudent to consider all contacts, beyond well-identified hydrogen bonds or salt bridges for example. It is especially true of highly polar interfaces such as the SARS-CoV RBDs vs. ACE2.

Using a 4A distance cutoff overall, the ACE2 - SARS-CoV-2 interface (PDB 6M0J) is formed by two <u>complementary surfaces</u> (Link 11), composed of mostly polar residues with 18 residues on the RBD (only one Arginine outside of the RBM) containing **6 aromatic** residues, 3 non-polar 2 Aromatic, 3Gly, 2 charged 1+/1-aromatic and polar vs. 20 residues on ACE2 with 3 non-polar, 1 Gly, 9 charged 4+/5- and **3 aromatic.** The interface with SARS-CoV-1 is compared in Figure 2. It also involves 20 residues on ACE2 with 16/20 being the same, but not necessarily the same interactions. A surprising number of aromatics are involved in conserved interactions (see later), specially conserved aromatics in SARS-CoV-1 vs SARS-CoV-2 RBMs (and RBDs overall). The <u>ACE2-SARS-CoV-2 interface (PDB 6M0J)</u> (Link 12) is composed of 12 H-bonds (green), 2 Salt Bridges (cyan) for a total of 36 non bonded contacts (black) with a cutoff distance of 4A, most of them polar (H-bonds calling is using both distance and angles criteria). The <u>ACE2-SARS-CoV-1 interface (PDB 2AJF)</u> (Link 13) is composed of 4HB, 1SB, 1CP, 1PP for a total of 34 nonbonded interactions

The ACE2-SARS-CoV-1 interface shows significant differences in the non bonded interactions “wiring” that can be schematically visualized in Figure 2. One should bear in mind that using a different experimental structure minute details may vary these numbers and that it is indeed important to analyze interfaces from alternate structures. For example, using PDB 6LZG <u>the SARS-CoV-2 the interface is composed of 7HBs, 2SBs but also 2 cation-Pi interactions for a total of 40 non bonded interactions</u> (Link 14). Using PDB 3SCI <u>the SARS-CoV-1 interface is composed of 5 HBs, 2SBs, 1 CP, 1 PP interactions for a total of 35 nonbonded interactions</u> (Link 15). Overall the interfaces contain between 34 and 40 non bonded mostly polar interactions. The devil is in the details however on how to call key interactions. We have more work to do in that respect. Overall these variations from identical protein structures solved independently show that interfaces are highly complex and require a great deal of attention to detail, one structure may identify a key interaction while one may miss it in using rigid distance criterion. Ideally, simulations should be performed to better understand molecular interactions and differences in the ACE2 binding interface with SARS-CoV-2 and SARS-CoV-1.

Interface analysis in literature - for example ^20^ << do we have a couple that mentions different residues?

Figure 2 compares interaction networks for these two viruses RBDs in binding ACE2. Conserved residues are represented in Red, vs. blue in the RBM for SARS-CoV-1 and SARS-CoV-2. ACE2 N terminal helical domain is in magenta and its other binding subdomain is in orange. Numberings are varying between structures and we used systematically in iCn3D, mapping to PDB numbers are available however through the links, and we provide mapping in Figure 4 for some key residues analyzed here. Hence one can notice immediately, despite various numberings! Conserved residues are in red **Y Y N Y G T G Y** and although many details can be analyzed through the links, we can here focus on identifying one conserved interaction in particular from one of these 8 conserved residues. First, notice that 4 of these are TYRosines! We focus on i.e. **Y453/Y440** in SARS-CoV-2/SARS-CoV-1 (Y135/Y118 in Figure 2) **and its conserved interaction with HIS34** (H16) **of ACE2** and will look hereafter at the rewiring of polar interactions around that **TYR-HIS** axis, located in the middle of the elongated ACE2-RBM interface.

**Figure 3.**
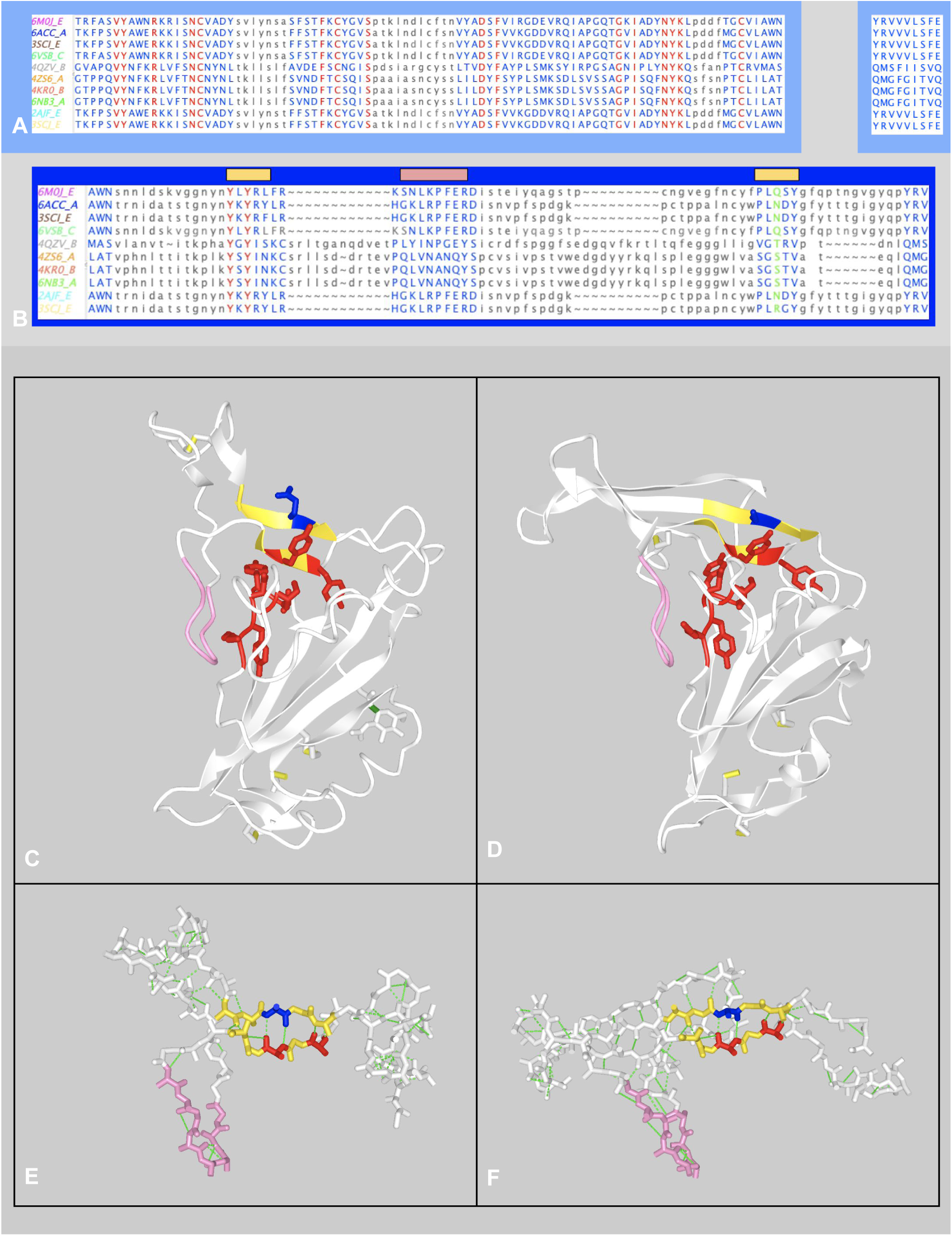
**A) Structural alignment of Beta Coronaviruses RBDs:** First Row the beta-sheet Framework (SkyBlue background). Second row: the plugin RBM (Dark blue background). A broader alignment using in addition OC43, HKU1, and MHV coronaviruses is available in Figures S1 and S2 **B) The RBM plugin contains a structurally conserved supersecondary structure (SSS)** comprising a central 2-stranded 5-residues long antiparallel beta-sheet (gold) and a flanking loop (pink). **C-D)** Visualization of the alignment in iCn3D of <u>highlighting of the conserved motif</u> (Link 16) in the structurally conserved antiparallel double-strand **YLYRL**//**PLQSY** in the RBM of SARS-CoV-2 where the residues Y135(453) and Q175(493) are themselves central, forming canonical backbone hydrogen-bonds. Also highlighted are the conserved **V**Y**…I…N**YK motif in the RBD framework that supports it. The structure-based multiple sequence alignment of the RBD/RBM from A) is loaded in Fasta format. Selected conserved residues are displayed in red. The structurally conserved antiparallel double strand of the RBM (gold) is interrupted by idiosyncratic structural elements, another structurally conserved loop between SARS and MERS beta coronaviruses flanks the RBD on the side (pink). Ninety-one backbone H-bonds give the SARS-CoV-1/2 RBM a quite rigid supersecondary structure, apart from the loop containing a Cys bridge when comparing and SARS-CoV-2. On MERS, an insertion extends the supersecondary structure (SSS) of SARS to 4 strands. **E-F)** The <u>RBM</u> <u>supersecondary structure</u> (Link 17) can be seen overall as a long doubly twisted two strand sheet, with a central two stranded-5 residues long beta-sheet. That SSS is common to SARS and MERS beta coronaviruses, colored in gold with the **YxY**135 (453) motif in red H-bonded at the backbone level to **Q**175 (493) in SARS-CoV-2 (see Figure 4 for details). In links, we superimpose both the conserved structural patterns (gold and pink) within 0.9A RMS. The structural superimposition of the central 2-strands aligns within 0.3A RMS between SARS-CoV-1/2. It aligns within 0.6A between SARS-CoV-2 and MERS (using Carbon Alphas). When <u>Comparing MERS to SARS RBM</u> (Link 18) the 2-central strands SSS, including the sequence pattern **YxY**135/138 motif is conserved, but also a very idiosyncratic loop (res. 141-148 SNLKPFERD in SARS-CoV-1/2 in pink) flanking the RBD framework on the side, structurally conserved but with no sequence conservation between SARS and MERS.

**Figure 4.**
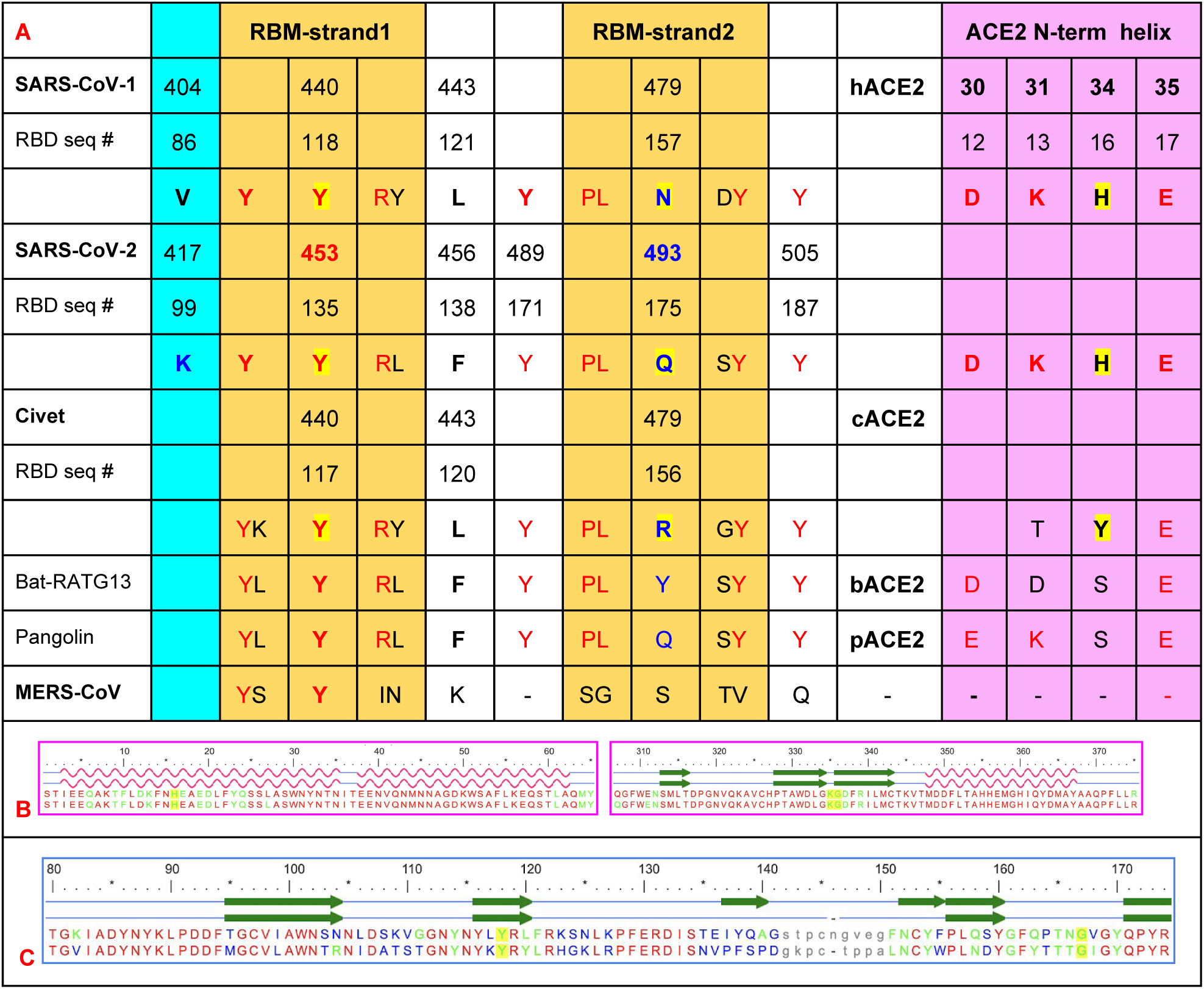
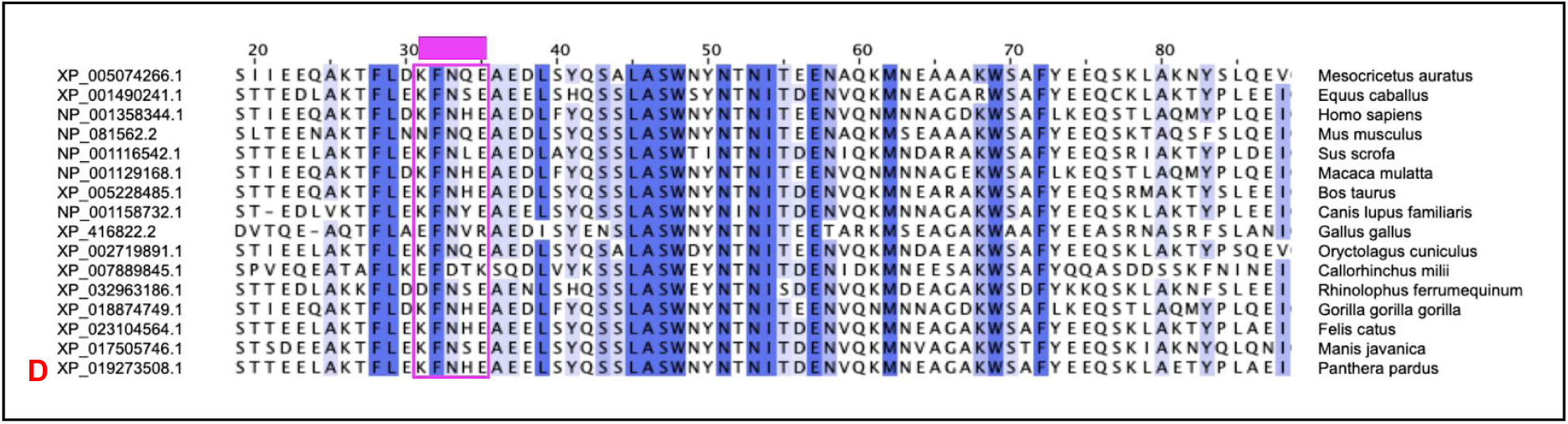
Conservation of amino acid residues in the central superstructural pattern of the RBM (Link 22). A) Comparison of SARS-CoV-2 selected residues: Y135/453 Yx**Y**Rx is at the center of the first strand, H-bonded with a second antiparallel strand PL**Q**xY to form a structurally conserved 2-strand central supersecondary structure (colored gold with Y in red (conserved) and Q in blue (not conserved). In Magenta the ACE2 main residues in contact and can vary, enabling rewiring to strengthen or weaken RBD/RBM to ACE2 binding interactions. The SARS-CoV-2 has an extra salt bridge indicated using a residue in the RBD framework, not the RBM (in cyan). Corresponding residues from SARS-CoV-1 and some animal sequences are indicated, as well as the corresponding residues in MERS for reference, although the latter does not bind ACE2, but DPP4. **<u>B-C</u>**) <u>Structure based</u> **<u>s</u>**<u>equence alignment of the quaternary complex RBD (A) -ACE2 (B) comparing SARS-CoV-1 and -2</u> (PDB 6M0J/2AJF) (Link 23) (red aligned/conserved and blue not conserved) with interface residues, conserved or not in green. Notice that most positions in either the RBM but also in the ACE2 RBS are conserved even if the residues are mutated. **<u>The interaction network rewires accordingly</u>**. Additional charged interaction involving K (green) (see Figure 2 - Link 10). **D) ACE2 sequences of selected mammals. Highlight of the 31-35 helical turn.** Sequence wise,in SARS-CoV-2 the central **TYR135/453** in the RBM is binding to **HIS16/34** in human ACE2, and that interaction is conserved in SARS-CoV-1. Interestingly cats, cows, and monkeys share the ACE2 K-H-E motif in positions 31-34-35 while it differs in bats, civets and pangolins, mouse, and dog. K13/31 before and E17/35 after H34 have different roles depending on the binding virus strain (see following paragraph). In A) we can see the corresponding sequences, and Figure 5 and the corresponding links describe differences. We also added a Shark Sequence. **(*)** Horseshoe Bat ACE2 sequence.

We will zoom in and compare a local subnetwork between SARS-CoV-2 vs SARS-CoV-1 vs. human ACE2 (Figure 4) in the following, but before we do so, **let’s look at at the sequence-structure conservation at a broader scale across beta coronaviruses to put this TYR453-HIS34 axis in an evolutionary context.**

#### Sequence and Structure conservation of beta coronaviruses RBD

Structures of known coronaviruses infecting humans, alpha and beta, are available (Table 1), alone or in interaction with their human cell surface receptors, and a comparative structural analysis gives an understanding of their structural and conformational architecture and evolution. MERS and SARS beta-coronaviruses genomes are available across species and can be analyzed in a structural context. Finally, a growing number of SARS-COV-2 viral genomes are being sequenced during the COVID-19 pandemic (https://www.gisaid.org/), and bringing sequence and possibly clinical information in a structural context may help foster a rational design approach of vaccines or of drugs.

**Table 1.** Known Structures of 7 known coronaviruses RBDs. Beta coronaviruses share the same RBD framework (Figures 2 A, B and 3A), and vary the RBM plugin (see Figure 2 A, B, and 3B) (see text). Figure 3A, B shows sequence variability for MERS and SARS beta coronaviruses. OC43 and HKU1 share the same RBD framework, their RBMs (not shown) have larger and more complex insertions (see Supplement Figure).

| CoronaVirus | Type | Receptor | Binding Domain | RBM | PDBid |  |
| --- | --- | --- | --- | --- | --- | --- |
| 229E | Alpha | N-amino peptidase | Barrel RBD | N/A | 6ATK | 21 |
| NL63 | Alpha | ACE2 | Barrel RBD | N/A | 3KBH | 22 |
| OC43 | Beta | unknown | RBD/RBM | Complex/Cys | 6NZK | 23 |
| HKU1 | Beta | unknown | RBD/RBM | Complex/Cys | 5GNB<br>5I08 | 24<br>25 |
| **MERS-CoV** | Beta | **DPP4** | RBD/RBM | 4-strands | 4KR0 | 26 |
| **SARS-CoV-1** | Beta | **ACE2** | RBD/RBM | 2-strands | 2AJF<br>3SCI | 27<br>28 |
| **SARS-CoV-2** | Beta | **ACE2** | RBD/RBM | 2-strands | 6M0J<br>6VW1 | 20<br>29 |

We look at a multiple structure alignment performed with the Cn3D ^30^ program, iCn3D predecessor, that allows highly accurate interactive alignment of protein structures, that automatic alignments cannot perform to that same level. RBD domains from SARS-CoV-1, SARS-CoV-2, and other beta coronaviruses from MERS were used to identify structural conservation patterns. As we observed in the architectural analysis, the RBD framework is highly conserved, and one can see in Figure 3a (blue background) a highly conserved framework with significant sequence conservation of key residues controlling its folding (with remarkably no sequence insertion). The RBM plug-in, responsible for binding to the receptor, however, shows extremely low sequence conservation, which shows its ability to adapt to different receptor targets.

One can observe a <u>significant structural variability in the RBM between MERS and SARS viruses</u> (Link 19), but only to notice that despite the extreme sequence variability, there is a level of structure resilience of the RBM of beta coronaviruses, beyond SARS strains, and there is one and only one conserved sequence pattern YxY (residues Y451 and Y453 in SARS-CoV-2) (Figure 3). The RBM SSS is <u>structurally conserved between SARS-CoV-1 and SARS-CoV-2 with an RMS of 0.304 A</u> (Link 20). The surprise is that it is <u>structurally conserved vs. MERS RBM, where is aligns within 0.64 A RMS</u> (Link 21) despite no sequence identity other than the **YxY** pattern and a significant insertion extending the beta-sheet

#### Focusing on the RBM supersecondary structure (SSS) and Y453-H34 centered binding

If we look at the alignment, it appears that the RBM subdomain (residues 117-191/435-509 in SARS-CoV-2) contains a structurally conserved supersecondary structure (SSS). Overall, the RBM plugin can be seen as an elongated domain sitting on top of the RBD framework. The idiosyncratic supersecondary structure is an irregular antiparallel 2 stranded 5 residue long beta-sheet with a first strand containing a conserved sequence motif Yx**Y**Rx (**strand1**) H-bonded to a second strand PL**Q/N**xY (**strand2**) in antiparallel. The central TYR135/453 sits on top of a conserved central folding pattern VY…I…NYK of the RBD framework (see Figure 3). Surprisingly, <u>The 2×5 residue long structural pattern with the YxY sequence motif is conserved in MERS</u> (Link 22) (see Figure 3), but has evolved two extra strands beyond that motif to bind DPP4 instead of ACE2. What is even more surprising is that conservation extends to OC43, HKU1, and MHH beta coronaviruses (see Figure S1), with larger RBM insertions. This conserved RBM supersecondary structure (SSS) is central and so well conserved structurally in itself and in its anchoring on the RBD framework that one can align entire RBDs simply by superimposing that central RBM SSS. The structural superimposition of the central 2-strands aligns within 0.3A RMS between SARS-CoV-½, it aligns within 0.6A between SARS-CoV-2 and MERS, and 0.9A RMS using both strands and the flanking loop together (using only Carbon Alphas). Incidentally, the structural pattern of the flanking loop is not uncommon in proteins; it has been described as a “GD box” ^31^. It is found, for example is the highly characteristic AB loop in IgV domains; it is also found in single-stranded left-handed beta helices, to which this flanking substructure bears some resemblance. Its sequence (res.143-148) is L**KPFE**R in SARS-CoV-2, (in pink in Figure 3), V**NANQ**Y in MERS-CoV (see Figure 3), V**GIGE**H in OC43, V**GVGE**H in HKU1, and should be V**NVGD**H for MHV after our realignment (see Figures S1 and S2).

Beyond folding, this RBM-SSS has a prominent role in binding: In SARS-CoV-2 **Y135** binds **H16** on ACE2, while **Q175 binds K13 and E17** that precede and follow H16 on ACE2. SARS-CoV-1 and SARS-CoV-2 share Y135/Y118 PDB #s Y453/440)yet differ with an essential **mutation Q175>N157**, a position that has been shown to reduce binding by a factor 30 if it is mutated to a Lysine (N479K) on hTor02 SARS-CoV-1 strain ^32,33^.

**In the following, we focus on this central 2-stranded antiparallel beta-sheet pattern** in SARS-CoV-1 and 2 in parallel with their Civets’ homolog that was thought to be a possible animal relay of SARS-CoV-1, for which we have a structure, as well as the pangolin and RATG13 Bat sequences that have been incriminated as the possible animal origin of SARS-CoV-2 ^34^ (see Figure 4).

#### ACE2 conservation across species and human polymorphism

##### Sequence-structure interface rewiring in SARS Coronaviruses

Numerous studies have been published analyzing various animal virus strains with human SARS-CoV-1 and SARS-CoV-2, especially in the RBD region that binds ACE2. It is not our aim in this paper to perform a review. For the sake of illustration, we compare interactions between various viral strains for which we have a structure (first 3 in Figure 4A), If we follow the central TYR135 (PDB# Y453 in SARS-CoV-2) residue that contacts the ACE2 HIS16 (PDB# H34), it binds HIS16/34 on ACE2, which is conserved across some species: human, monkeys, cats, cow, while it differs in dog, mouse, civet, Bat RATG13, and pangolin ^34^ (see Figure 4).

##### Comparative binding in the RBD-ACE2 interface between Civet and Human viruses

Palm civet was thought to be the animal origin of SARS-CoV-1, and it was believed that the major species barriers between humans and civets for SARS infections are the specific interactions between a defined receptor-binding domain (RBD) its host receptor, angiotensin-converting enzyme 2 (ACE2) ^33^. The structure of a chimeric human-civet ACE2 receptor with the critical N-terminal helix from civet shows a conserved interface ^28,32^. If we compare the ACE2-RBD interfaces of SARS-CoV-1 and SARS-CoV-2 but also the ACE2 chimera vs. the civet’s RBD <u>(6M0J vs. 3D0I), again the central RBM-ACE2 interface is conserved overall</u> (Link 24), we can notice a number of subtle changes vs. the RBD structures of SARS-CoV-1 SARS-CoV-2, especially with the RBM central region interface to ACE2 N-terminal helix we have been outlining (Figure 3-4). In fact, it has been shown that “a major species barrier between humans and civets for SARS-CoV-1 infections is the adaptation of residue 479 on RBD”. It corresponds to Q175/493 in SARS-CoV-2, (PDB: 6M0J) vs. N157/479 in SARS-CoV-1 (2AJF) interacting with residues K31 and E35 in hACE2 (K13/E17), corresponding to T31 and E35 in cACE2 (T13/E17).

##### Mapping human ACE2 polymorphism on structure

We have analyzed the interface between the virus RBD and ACE2. We have looked at mutations in the RBD between different viruses SARS-COV2 vs SARS-COV1 vs. MERS. Yet the ACE2 also varies from patient to patient. Since mutations at the ACE2 interface are important to modulate binding, the question one may ask is what would be the impact of human ACE2 polymorphism in binding to SARS-CoV-2 and the possible variations in pathogenicity leading to COVID-19. iCn3D accesses the human SNP database seamlessly, and one can look at ACE2 polymorphisms in a structural context. In Figure 5, we can visualize the interacting residue network with SNP positions marked (in green) that can affect the binding.

**Figure 5.**
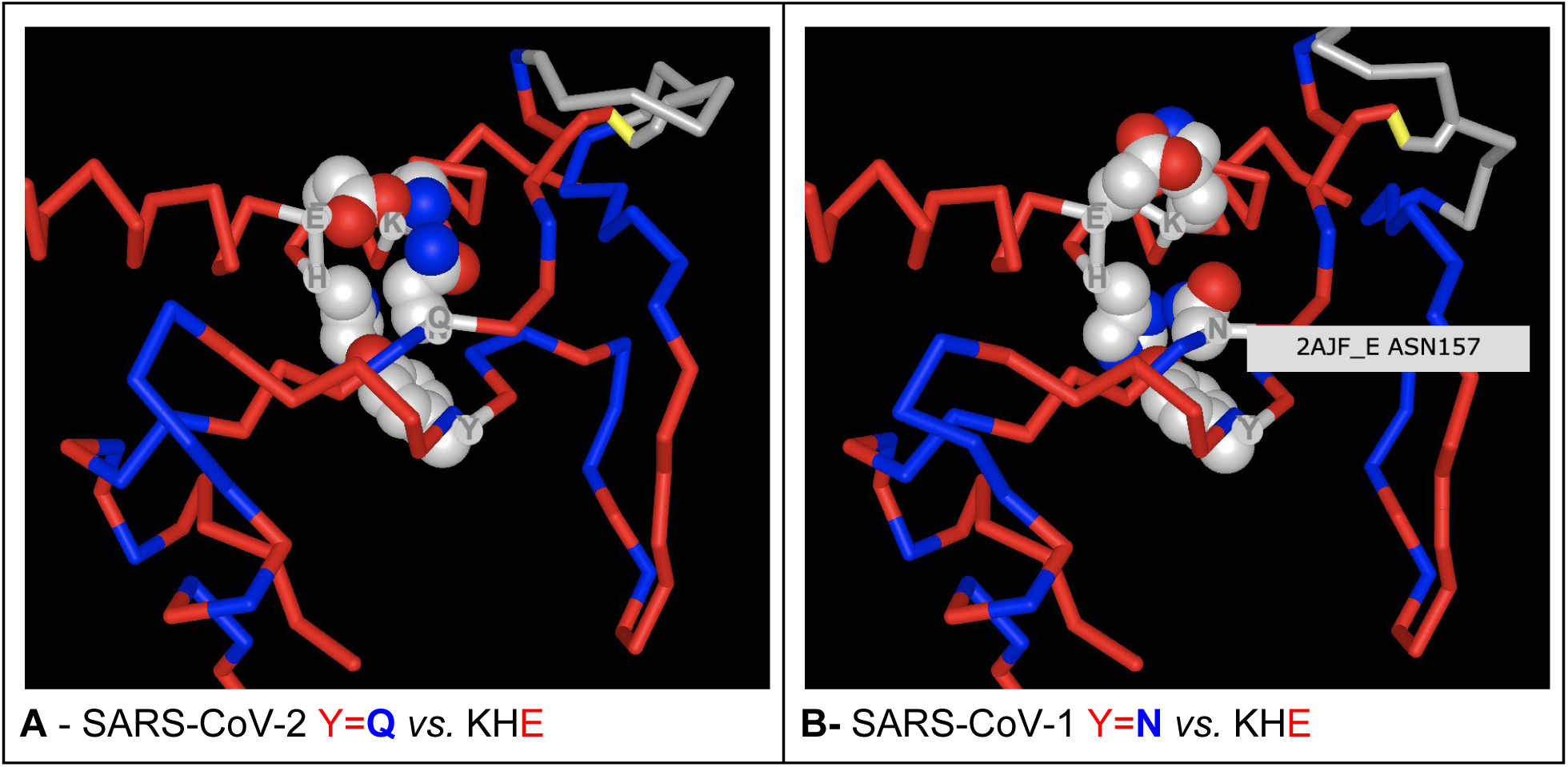
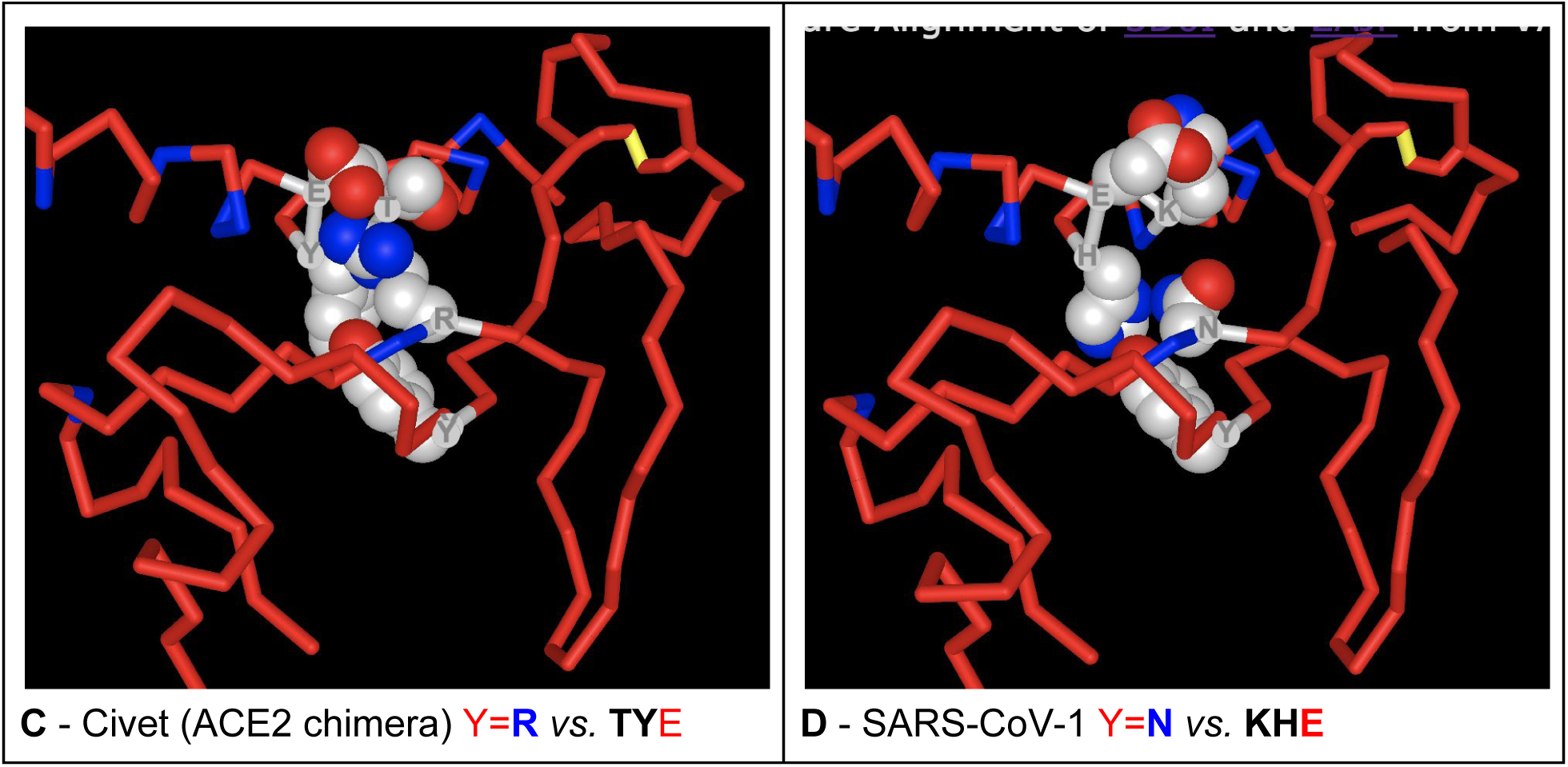
A-B) Comparison of the SARS-CoV-2 vs. SARS-CoV-1 binding of the RBM-SSS to hACE2 (Link 25). While Y135/453 in strand 1 of the central SSS binds ACE2 H16/34, in SARS-CoV-2 the residue in vis a vis in strand 2 Q175/493 binds simultaneously K31 and E37 through a hydrogen bond (see Figure 2). while in SARS-CoV-1 where Q>N at that position (N479), the shorter side chain does not reach as far out to contact K31-E37 that form an internal salt bridge within ACE2. The comparison is made easy using the alternate command in the iCn3D link one can see (“a” letter on keyboard) to alternate between the 2 homologs superimposed. **C-D)** <u>Comparison of the Civet-CoV/cACE2 (chimera N-terminus) vs. SARS-CoV-2/hACE2</u> (Link 26). The Civet’s strain (cGd05 - PDB 3D0I) harbors an Arginine residue R479 at the position occupied by an Asparagine residue in SARS-CoV-1. In binding to Civet’s ACE2 where threonine is at position T31 (while HIS16/34 is mutated to a TYR Y34) a salt bridge is formed between R479 and ACE2 E37. In human ACE2, where position 31 is a Lysine K31, it has been shown that a single N479K or N479R mutation on human viral RBD decreases the protein’s affinity for hACE2 by over 30-fold 32, which is understandable for a complex with two positively charged residues. iCn3D does not perform simulations, but visualization and structural analysis give ideas on what computational properties may be explored through simulation programs. For example, in this case, having identified the pattern, we assessed the difference in binding by computing the relative binding free energy using a free energy perturbation (FEP) simulation scheme, obtaining a ΔΔG = 6.3 (N479K) and 9.1 (N449R) Kcal/mol respectively. In SARS-CoV-2 however, the mutation is N479Q wrt SARS-CoV-1 on the contrary strengthens the binding. The FEP analysis for this mutation results in a favorable ΔΔG = -3.0 Kcal/mol (see Supplementary information for details)

**Figure 5.**
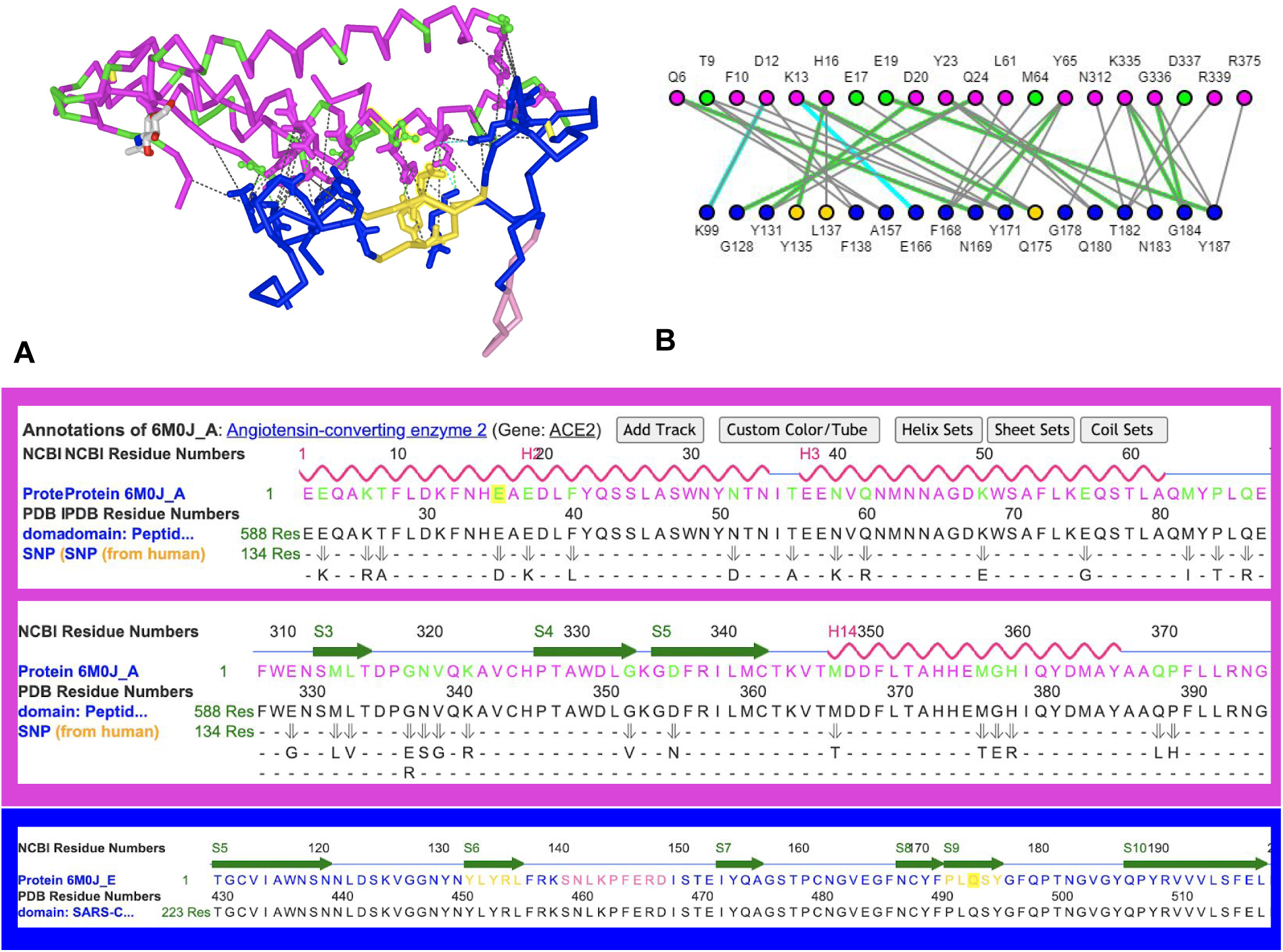
ACE2 human polymorphism at the RBD interface (Link 27). **A) ACE2 is colored in magenta, and in** green are positions **within ACE2 subdomains that** can vary (dbSNP). In blue, the RBD/RBM contacts ACE2 and in gold residues within the strand1-strand2 in the RBM. For example E17D or more likely E19K would affect the interface in the details of binding the RBD/RBM. **B) the corresponding network of interactions** (see Figure 2 for details).

As we have seen at the beginning, <u>the binding between the RBD and ACE2</u> (Link 10), concentrated in the RBM (but for one residue K99 in the RBD framework - see Figure 2) forms an extensive elongated binding interface composed of c.a. twenty residues on both sides and ca. 40 residue-residue interactions (see the paragraph **comparing networks of interacting residues in SARS-CoV-1 and SARS-CoV-2 *vs.* ACE2).** We have only focused on a central region involving the RBM-SSS that we have outlined using the same token highlighting some of the analytical capabilities of iCn3D that allowed us to investigate structures and sequences in parallel, focusing on molecular interactions that enable viral entry. Whether TYR453 and the central RBM-SSS may be specifically targeted by cross-neutralizing antibodies, or even small molecules, need further investigation. Neutralizing antibodies isolated from recovered patients such as <u>b38 cover the entire RBM of SARS-CoV-2</u> (Link 28) ^35^. Deep insights can come from comparisons in binding between viruses, their receptors, and neutralizing antibodies. We leave to readers the opportunity to find other interesting “key” interactions using iCn3D, with the links provided as starting points. They can be used directly, or as templates to study new structures as they become available.

#### Epilogue: The necessary interplay sequence-structure analysis

Structure alignment can guide the proper comparison of sequences, significantly improving the chances to discover hidden patterns. This has been well known, but correlating 1D and 3D data involves specialized techniques that have been difficult to use for those not trained in their use. This type of integrated analysis is now accessible through iCn3D, greatly facilitating this work, as shown in the previous examples. iCn3D and the seamlessly integrated remote database infrastructure it benefits from makes the job more intuitive, and the actions are better integrated, as we have shown in the previous examples applied to Covid-19 relevant problems.

Different viruses are frequently compared through sequence alignment, for instance while attempting to characterize RBDs. However, sequence-based alignments can be misleading, for example, in some highly evolved regions (see Supplement Figure S2 comparing sequence alignment in ^36^ Figure 1, mispositioning an insertion in MERS vs. SARS-CoV-2/1, vs. Figure 3.B and <u>Link 19</u> (Table 2). Therefore, it is critical to get accurate alignments if the aim is to identify viral attachment sites targeted by inhibitors or as vaccine epitopes, possibly common to various virus strains. Protein immunogens, in particular, need to be conformationally correct ^3^. Our analysis reveals that betacoronavirus (SARS-CoV-1 vs. SARS-CoV-2 vs. MERS) display a small common sequence-structure pattern with remarkable structural stability that can be used as the focus for structure-based design efforts.

**Table 2.** Summary List of iCn3D Share_Links with the corresponding text. The corresponding “gallery” is also added as a supplementary HTML file S2. A gallery is generated by clicking the links and saving them with the option “iCn3D PNG Image”, and then concatenating the saved HTML files.

| Link # | Text | Share Link |
| --- | --- | --- |
| 1 | the SARS-COV-2 Receptor Binding domain interacting with the ACE2 receptor | https://structure.ncbi.nlm.nih.gov/icn3d/share.html?6soVn4ka47phy7a66 |
| 2 | antibodies against SARS-COV-1 vs. SARS-COV-2 | https://structure.ncbi.nlm.nih.gov/icn3d/share.html?bkYBrK8WcyM8kD8M6 |
| 3 | mutations across coronaviruses: beta coronaviruses sequence alignment is visualized in iCn3D (CDD CD21470) | https://structure.ncbi.nlm.nih.gov/icn3d/share.html?ZuAQgrZHXkNoc1q37 |
| 4 | domain composition in ribbon representation | https://structure.ncbi.nlm.nih.gov/icn3d/share.html?uNWzXjGCn379S3Gu5 |
| 5 | SARS-COV-1 spike is highly homologous and can be superimposed for the whole trimer (PDB 6CS2) | https://structure.ncbi.nlm.nih.gov/icn3d/share.html?vRWRTWJGMR9j5XHt5 |
| 6 | the spike Trimer structure (6CS2) of SARS-COV-1 binding the full length ACE2 peptidase (PDB 6M17) | https://structure.ncbi.nlm.nih.gov/icn3d/share.html?o4TZdnKVd6JcPw8z8 |
| 7 | compare SARS-CoV-1 and SARS-CoV-2 RBD interfaces by superimposing the two complexes (quaternary structures) 6M0J vs. 2AJF | https://structure.ncbi.nlm.nih.gov/icn3d/share.html?AMyajRGMPjY2HAu3A |
| 8 | compare SARS-CoV-1 and SARS-CoV-2 RBD interfaces by superimposing the two complexes (quaternary structures) 6M0J vs. 3SCI | https://structure.ncbi.nlm.nih.gov/icn3d/share.html?qNgcsZQARcxFrcSR9 |
| 9 | side by side comparison of SARS-CoV-1 and SARS-CoV-2 RBDs | https://structure.ncbi.nlm.nih.gov/icn3d/share.html?xwDbyM9eGSCzS57d6 |
| 10 | SARS-CoV-1 - ACE2 vs. the SARS-Cov-2 - ACE2 | https://structure.ncbi.nlm.nih.gov/icn3d/share.html?UjBF9jtDEUPyKXHo9 |
| 11 | the ACE2 - SARS-CoV-2 interface (PDB 6M0J) is formed by two complementary surfaces | https://structure.ncbi.nlm.nih.gov/icn3d/share.html?Qebj9RQymo49ryCy9 |
| 12 | ACE2-SARS-CoV-2 interface (PDB 6M0J) | https://structure.ncbi.nlm.nih.gov/icn3d/share.html?4T4qW9JJj9gw14cW8 |
| 13 | ACE2-SARS-CoV-1 interface (PDB 2AJF) | https://structure.ncbi.nlm.nih.gov/icn3d/share.html?HCAuQVN47ZkcJGMK7 |
| 14 | the SARS-CoV-2 the interface is composed of 7HBs, 2SBs but also 2 cation-π interactions for a total of 40 non bonded interactions | https://structure.ncbi.nlm.nih.gov/icn3d/share.html?Tx9U4g7DBPrA88aUA |
| 15 | the SARS-CoV-1 interface is composed of 5 HBs, 2SBs, 1 CP, 1 PP interactions for a total of 35 nonbonded interactions | https://structure.ncbi.nlm.nih.gov/icn3d/share.html?UsQVMtdGSRQ9qV319 |
| 16 | highlighting of the YxY conserved motif | https://structure.ncbi.nlm.nih.gov/icn3d/share.html?4GyBkQNwt5v5XqhB7 |
| 17 | RBM supersecondary structure | https://structure.ncbi.nlm.nih.gov/icn3d/share.html?BpW1vpX1M4vCSnrHA |
| 18 | Comparing MERS to SARS RBM | https://structure.ncbi.nlm.nih.gov/icn3d/share.html?42EwYyUPzBbpRe8U6 |
| 19 | significant structural variability in the RBM between MERS and SARS viruses | https://structure.ncbi.nlm.nih.gov/icn3d/share.html?m5syRCztf09C2fdX7 |
| 20 | structurally conserved between SARS-CoV-1 and SARS-CoV-2 with an RMS of 0.304 Å | https://structure.ncbi.nlm.nih.gov/icn3d/share.html?2e5yreQjVG7fuL8o7 |
| 21 | structurally conserved vs MERS RBM, where is aligns within 0.64 Å RMS | https://structure.ncbi.nlm.nih.gov/icn3d/share.html?3CXbv1AsUGdXTDTZ9 |
| 22 | The 2x5 residue long structural pattern with the YxY sequence motif is conserved in MERS | https://structure.ncbi.nlm.nih.gov/icn3d/share.html?D4cq5tDXS3GNyh51A |
| 23 | Structure based sequence alignment of the quaternary complex RBD (A) -ACE2 (B) comparing SARS-CoV-1 and -2 | https://structure.ncbi.nlm.nih.gov/icn3d/share.html?LWTxP49jjx5j7rit9 |
| 24 | (6M0J vs 3D0I), again the central RBM-ACE2 interface is conserved overall | https://structure.ncbi.nlm.nih.gov/icn3d/share.html?RFQSmr7BoafTUJF8A |
| 25 | Comparing SARS-CoV-2 vs. SARS-CoV-1 binding of the RBM-SSS to hACE2 | https://structure.ncbi.nlm.nih.gov/icn3d/share.html?YtjxMkvYqAcDsvXt7 |
| 26 | Comparing Civet-CoV/cACE2 (chimera N-terminus) vs. SARS-CoV-2/hACE2 | https://structure.ncbi.nlm.nih.gov/icn3d/share.html?K24q8LRgpyvK1RPw5 |
| 27 | ACE2 human polymorphism at the RBD interface | https://structure.ncbi.nlm.nih.gov/icn3d/share.html?BbfXuWEP4y9eCzNp9 |
| 28 | b38 cover the entire RBM of SARS-CoV-2 | https://structure.ncbi.nlm.nih.gov/icn3d/share.html?JidSY99aAwRngkkV6 |

While a sequence alignment may be guided by structure, the converse is also true. Tracing a sequence in an electron density map, when that map is not well resolved, can be a very challenging task. We extended our structural alignment in (Figure 3.A, B) to other beta coronaviruses presenting a third type, more complex RBM (Table 1), i.e., OC43 and HKU1 as well as MHV (Murine Hepatitis Virus) (Figure S1). We found that the previously described RBM supersecondary structure is still conserved in these viruses. Its structural resilience is remarkable in accommodating insertions that differentiate and bind diverse target receptors, some yet unknown. In performing this analysis, we found structures that could benefit from the study here presented to guide the refinement process (i.e., MHV ^37^ -PDB 3JCL-). The MHV structure, for example, introduces a shift in sequence while attempting to fit the electron density map, resulting in a local misalignment and, as a consequence, a likely misassigned Cys bridge.

A CDD evolutionary sequence alignment ^38^ of beta coronaviruses (<u>CD21470</u>) shows that sequences of a variety of viruses have homologous sequence patterns that can be used to guide sequence tracing in low-resolution electron density. In fact, the **Y135/453 conservation can be extended to OC43, HKU1, and MHV**, the pattern Yx**Y** in SARS-CoV-1, SARS-CoV-2, and MERS mutates to VV**Y**. In the case of MHV (PDB 3JCL), structural alignment of other structures confirms that homologous sequence patterns are also structurally conserved. Hence the sequence RBM-SSS-strand 1 can be realigned locally, and from there, strand 2 and the RBM-SSS flanking loop can also be followed. Another small sequence shift can be observed on the structure of HKU1 (PDB 5I08). These local sequence realignments are available in Figure S1, where it can be seen that all known beta coronaviruses exhibit the same supersecondary structure in the RBM with a highly conserved Tyrosine, as previously described.

In short, we took the problem of parallel characterization of the RBD domains, including diverse RBM subdomains across a variety of coronaviruses, paying special attention to SARS-CoV-1, SARS-CoV-2, and MERS-CoV, to see how far our methods could go to help current research and researchers, and how we should push them further in the future. Through these analyses, we observed **a common sequence-structure pattern across different viruses that have not been analyzed in-depth before**.

We could not have observed that pattern without a rigorous structure based sequence alignment across multiple coronaviruses and, reciprocally, a parallel analysis of diverse structures guided by evolutionary sequence alignments.

We have shown how structural analysis methods in iCn3D can be used in the context of COVID19 related structures exploration. Beyond the technological benefits of web-based visualization, web-based data and annotations sharing, and collaborative analysis capabilities, we demonstrate the scientific merit of performing a simultaneous sequence-structure analysis.

### 3. Collaborating, Data Sharing, and Publishing

#### Collaborative Research

Research collaborations can be facilitated by exchanging links. A link can be passed between scientists in a “**passive**” mode to visualize annotated structures, highlighting specific features, properties, interactions through multiple labeled representations. Collaboration can become **active**: a link is live, it is not just a simple visual representation, it contains the software that enables analysis. An analysis is passed at a given stage and it is in a given “state” that can be used for further analysis and edition, making iCn3D a collaborative diSARS-CoV-very tool.

#### Lifelong links

Lifelong links are currently hosted by NCBI. It could be hosted by publishers or some other data service. That hosting currently mediates Google Lifelong links in order to ensure continuity of access. By this mechanism it is now possible to update links and maintain them over time, as links addresses are managed. The solution to making links truly lifelong and maintainable is to give a permanent name, as we do for a domain name vs an IP address - one can host the same domain name on any server with any IP address. The same concept is applied to links for which NCBI acts as a Link registry and can be evolved towards using DOI (www.doi.org) scheme in the future. In all cases, links offer a way forward with perennity and are a game-changer to share information, methodologies, and data itself.

#### Data sharing through links

Where is the data? Currently, analytical (processed) data is spread as text, tables, figures, and supplemental files in published papers, while sequence or structural data is available in depositories such as the PDB or NCBI and others. A link is in fact a set of commands interpreted by iCn3D that access and process the data in a well-defined manner, making links, data, and entire analyses they encapsulate FAIR ^2^, i.e. **Findable, Accessible, Interoperable, and Reusable**. Reusability of a link, as mentioned before can be passive, but is also de facto **Extensible**, by design, and a key to research **Collaboration**. Any published link makes the data FAIR and makes an analysis **Reproducible.**

Data generated by analyses are computed on the fly at any instantiation of a link using basic data stored at NCBI and PDB. Analytic data is the product or primary data processed by an analysis method encoded in iCn3D. All commands producing numerical data such as tables of molecular properties or molecular interactions allow the export as files, hence instead of using supplemental data files in preprints of papers or attachments to emails, they are all intrinsically contained in a link. The concept of reproducibility is internalized in iCn3D, and data sharing is intrinsic to a link. In addition, in order to provide perennial accessibility, reproducibility, and reusability over long periods of time to analyses, data and annotations, iCn3D released versions are archived (https://www.ncbi.nlm.nih.gov/Structure/icn3d/icn3d.html#log). Should a future version change some features affecting a published link, that link can be replayed with the original iCn3D version that was used to generate it, based on its recorded creation date.

Hyperlinks were native to the WWW from its origin, so iCn3D Share Links are a natural evolution, using them to share molecular scenes, data, and annotations. Links can be served through a web server, currently through the NCBI server, but some other external scientific data services have recently started to use them, such as OPM (https://opm.phar.umich.edu/).

Down the road, links could be used as data and annotation aggregators, provided the original data location or computer servers producing data remain accessible. This is an issue beyond the current use and would require on-going coordination between data services, but it could be envisioned, especially at a time when data sharing has become mandatory for, for example, research funded by the NIH (https://grants.nih.gov/policy/sharing.htm).

#### Peer Review and Publication

Beyond discovery, links are designed to disseminate research results as data and annotations. They enable the **sharing** of annotated 3D structures for publication in preprints or peer-reviewed papers. Links are perfect peer to peer exchange vehicles as well. They could be invaluable to **peer review**, since they present researchers’ data and analysis, avoiding the use of supplement data and figure files that are currently needed by reviewers. Many supplement files may be eliminated from preprints and published papers through the use of web links. Another interesting aspect of links is the ability to cite data directly. A lifelong link is similar to a DOI and can be cited since it can be addressed from anywhere on the Internet. Current links are hosted by Google and are being made maintainable (editable) by NCBI. They could very well be hosted anywhere in the cloud and managed by individual organizations or publishers.

#### Education

Lastly, links are a perfect means to disseminate scientific knowledge, and teachers have been faster in picking up their value in education. iCn3D HTML galleries can represent a graphical summary for a paper, a course, or even a scientific presentation. Since it is an HTML file, it can simply be edited to provide a simple means to annotate and document a scientific project. For example, the Supplementary HTML file S2 summarizes the entire set of links (Table 2) of this paper graphically and we continuously append new links associated with new analyses in <u>the iCn3D gallery</u>.

## Future Developments

Numerous improvements can be envisioned, from adding analysis functionality to extending its use for group editing. iCn3D is open source to help foster a community of interest that can collaborate on future developments.

As we implement new functionalities, we will inevitably increase the program complexity. There are therefore different directions to consider for the future to reduce technical complexity. One is to use a viewer only version of the program for purely passive visualization use. A link may encapsulate a highly sophisticated analysis, yet there is no need of any particular knowledge to visualize molecular scenes. This “light” implementation may best support the rapid publication and dissemination of 3D annotations. Beyond visualization, in its interactive structural analysis mode, it becomes an active collaboration tool: a link is then not just an endpoint, it is a starting point for further exploration.

Another direction towards minimizing complexity in interactive mode, while increasing functionality is in increasing analysis automation, and possibly using AI, which may greatly benefit from the model of centralized repositories used by iCn3D. The evolution from interactive to automated analysis should open the door to making sophisticated analyses simple and accessible to non-experts. It is also a necessity for experts to be able to scale up and deal with large ensembles of structures rather than one or a handful at a time. We have currently no tool to deal with 150,000 structures available in the PDB database, or even a few hundred, while some structural families, such as antibodies, include thousands of structures. This opens a whole new research and development area in itself, beyond interactive analysis.

## Supporting information

Graphical Abstract of Links with active Links

## ACKNOWLEDGEMENTS

We thank the CDD group at NCBI, especially Aron Marchler-Bauer, James Song, Myra Derbyshire, Narmada Thanki, Noreen Gonzales, who provided us with the evolutionary alignments of beta coronaviruses, pointing us to the conservation of sequence patterns in OC43, HKU1 and MHV (see Figure S1 and S2).

On a personal note, PY would like to acknowledge the enduring inspiration that Cyrus Levinthal has exerted in the use of interactive structural analysis (and by extension modeling).

*This research was supported in part by the Intramural Research Program of the National Cancer Institute and the National Library of Medicine, NIH.*

## SUPPLEMENTS

**Figure S1.**
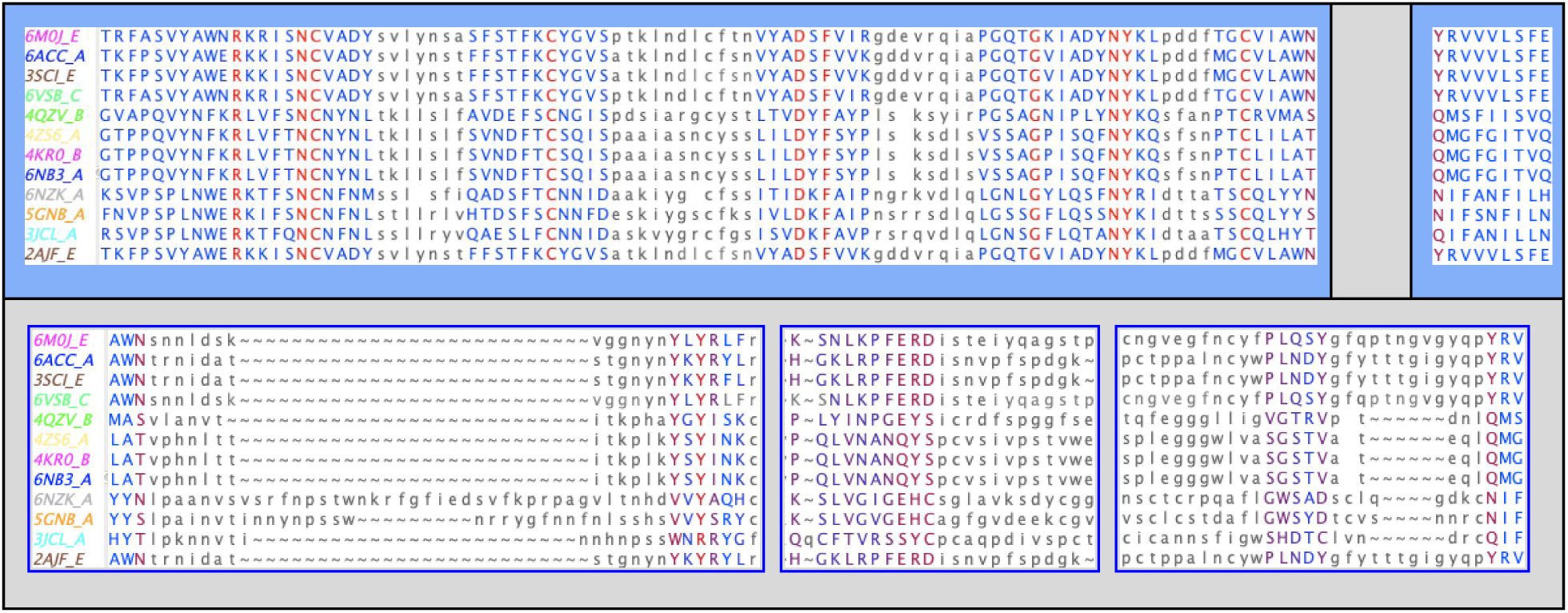
Structural alignment of known Beta Coronaviruses RBDs: Similar alignment to Figure 3A, with in addition **OC43, HKU1, and MHV**. For clarity we omit additional insertions in the RBM plugin vs. **SARS and MERS** coronaviruses. These insertions increase the RBM size and bring a significant structural complexity due to disulfide bridges. What is remarkable however, is the structural conservation of the RBM SSS despite significant insertions. The first Tyrosine Y133/451 in strand 1 is however, not as conserved as Y135/453, as in, for example, HKU1 or OC43 coronaviruses (PDBids 5GNB and 6NZK) where it is replaced by a Valine. This broader <u>coronaviruses sequence alignment is visualized in iCn3D</u> (Link 3). In the structure of a mouse hepatitis virus (MHV) (PDBid: 3JCL), the 3D superposition also shows a structurally conserved RBM strand 1 where the sequence **WNR** matches the **Y/VxY** pattern location, yet an evolutionary-based sequence alignment of beta coronaviruses (<u>CD21470</u>) finds a **VVY** match as in HKU1 and OC43, located in sequence right after the (PSS**WNR**RYGF) sequence conserved in OC43 and HKU1. The disagreement is most likely the result of the region being poorly defined in the electron density map, as stated by the authors of the structure and seen in the following figure (Figure S2) Whether an artefact of the EM structure or a sequence shift during evolution, the RBM structural pattern (antiparallel strand1-strand2-flanking loop) is conserved across all know beta coronaviruses.

**Figure S2.**
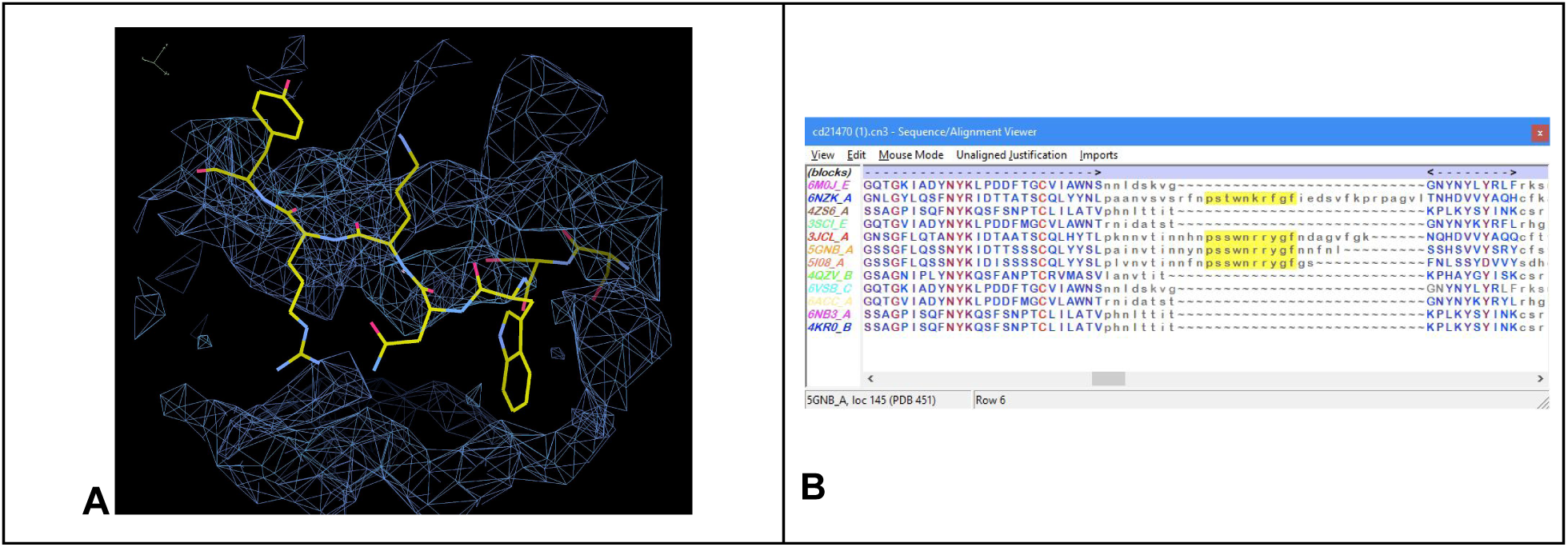
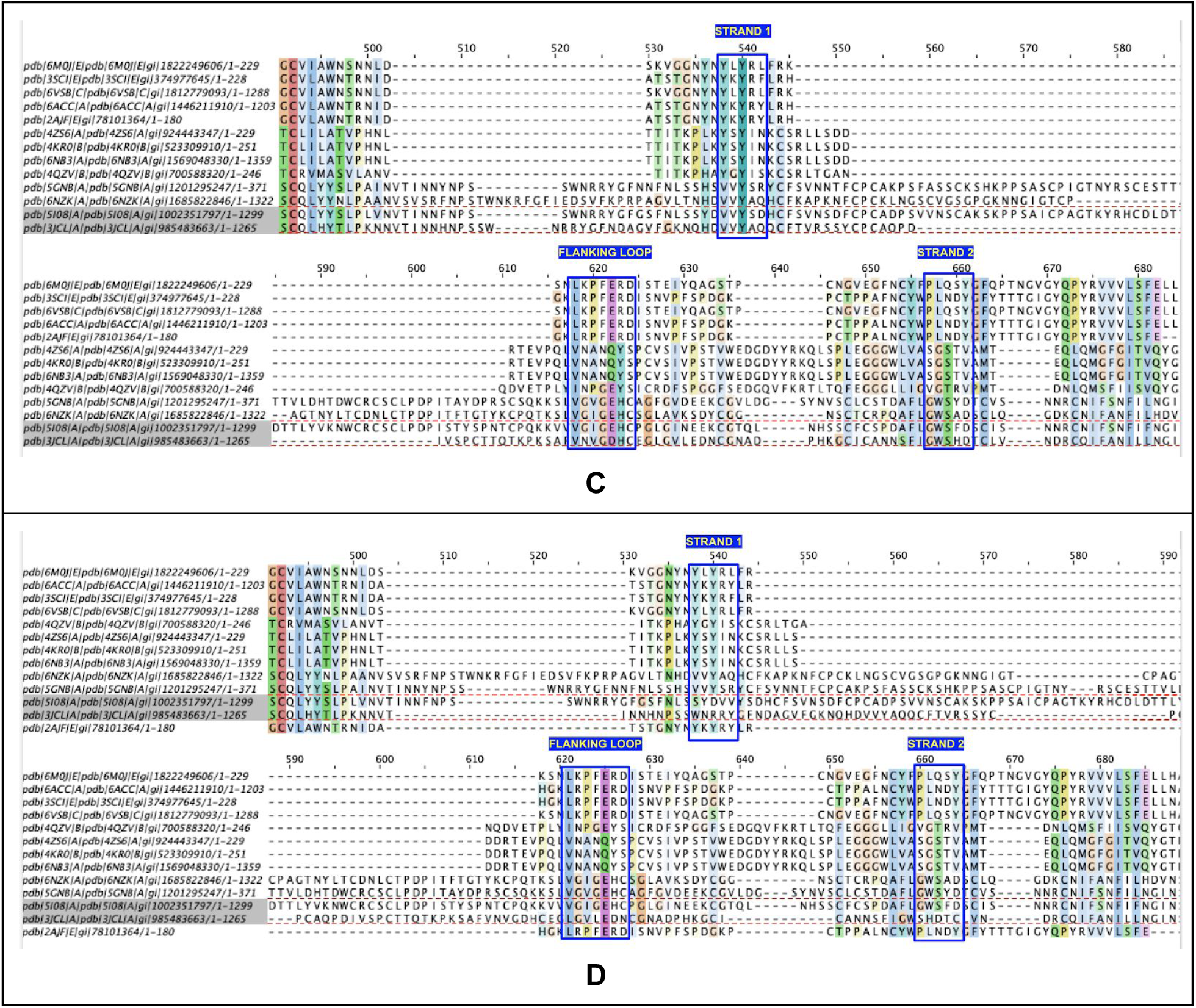
RBM subdomains of beta coronaviruses. **A) electron density of** MHV (Murine Leukemia Virus) structure 3JCL in the RBM region. The sequence **WNR** was fitted in place of VV**Y** **B) Beta Coronaviruses CDD (**<u>CD21470</u>) showing the conservation of the PSSWNRRYGF sequence in OC43 and HKU1 as well as MHV **C)-Optimized Sequence alignment of beta coronaviruses RBMs** taking into account homologous patterns observed in the CDD (B) where Strand 1 aligns Yx**Y** motif of SARS/MERS vs. VV**Y** for OC43/HKU1/HKU4/MHV. In Strand 2 similarly a GWS pattern matches in these four viruses the PLQ/N sequence in SARS. Similarly the GD-box like flanking loop, between strand 1 and strand 2 aligns GxxE between OC43/HKU1/HKU4/MHV **D)-Optimized structure alignment of beta coronaviruses RBMs** where in 3JCL WNR was fitted (see A) and aligns with other structures VVY in strand 1’, in strand 2 the structure presents a shift of 2 residues of GWS when compared to other structures C). The HKU1 (PDB 5I08) structure, in the same way shifts by 3 residues strand 1 matching VV**Y** with the preceding SYD residues, yet strand 2 and the flanking loop are well positioned.

**Figure S3.**
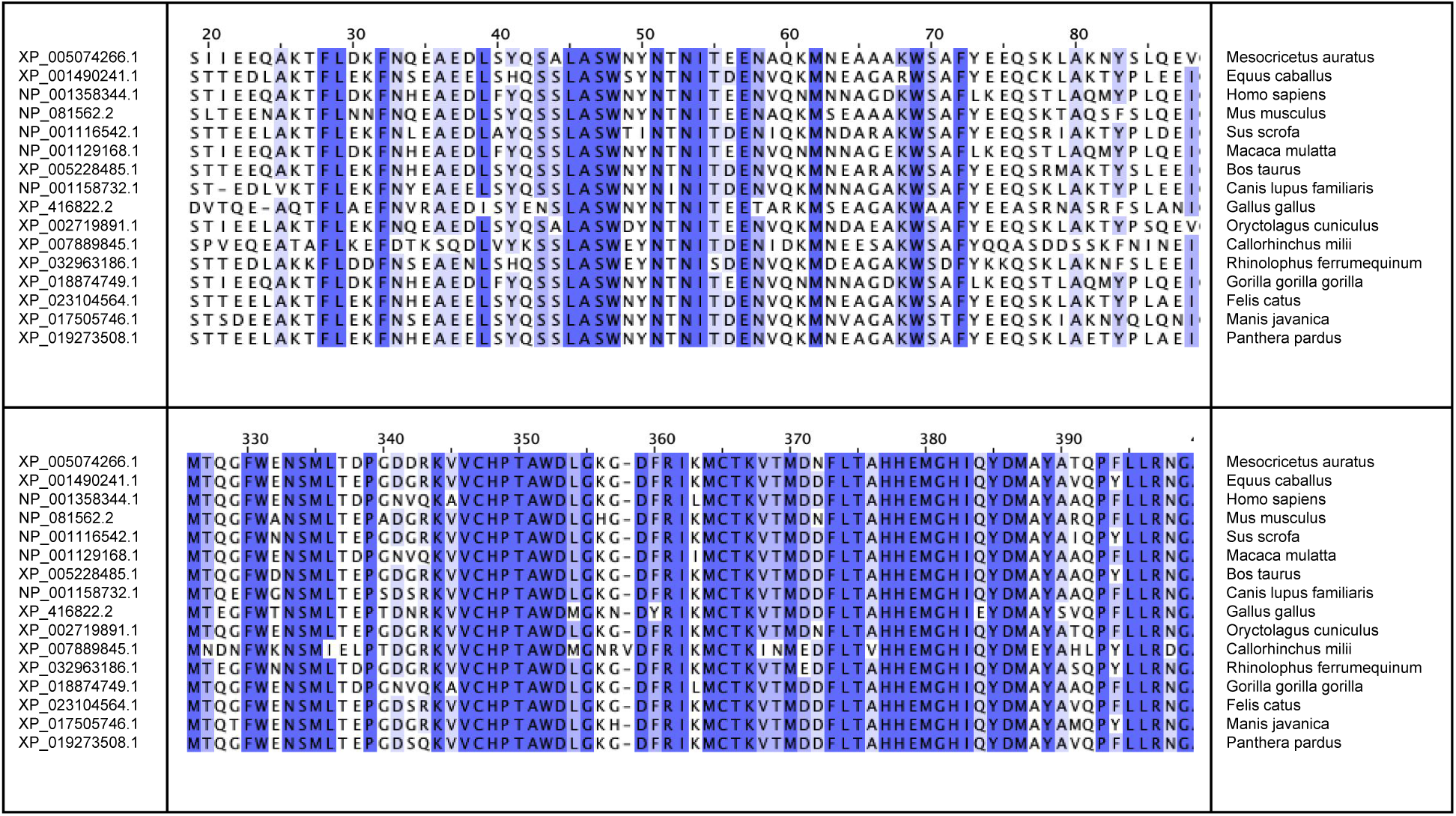
ACE2 sequences of selected mammals. Two subdomains contain the residues forming the receptor binding site (RBS) of ACE2, 1. N terminal domain (res 19-90) and 2. subdomain (res. 326-400) [see Figure 2-3-4] for more details.

### Free energy calculations

ΔΔG values were computed using the Free Energy Perturbation slow growth calculation approach as described in [1] using Gromacs 5.1.4 [2] and the PMX package [3] considering the “unbound” geometries (instead of the “unfolded” states as described in [1]) The ΔΔG values were computed in the forward and backwards direction. The values reported are the average of the two. The Amber99sb force field was used for all FEP calculations. Initial coordinates were obtained from the RCSB and preprocessed using Yasara version 20 [4] and the “Clean All” macro using the NOVA2 force field, followed by an energy minimization using the same force field and default parameters using the Options / Energy Minimization macro. The coordinates were then subjected to the regular relaxation in a solvent bath as described in [1] before the molecular dynamics trajectories were collected.

<u>[1]</u> https://www3.mpibpc.mpg.de/groups/de_groot/cecam2015/peptide_mutation/

<u>[2]</u> http://manual.gromacs.org/documentation/5.1.4/

<u>[3]</u> https://github.com/dseeliger/pmx/

<u>[4]</u> http://www.yasara.org/

