## Supplementary material for "Using iCn3D and the World Wide Web for structure-based collaborative research: Analyzing molecular interactions at the root of COVID-19": Graphical Abstract of Links with active Links

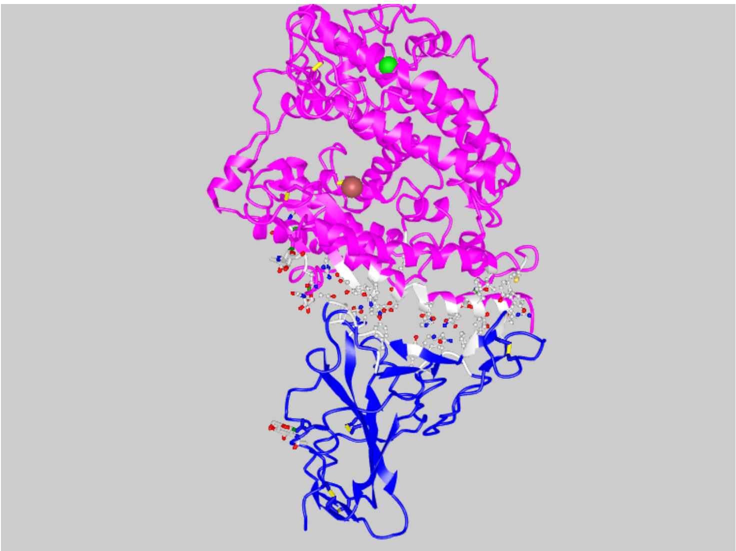

Link 1: PDB 6M0J: the SARS-COV-2 Receptor Binding domain interacting with the ACE2 receptor

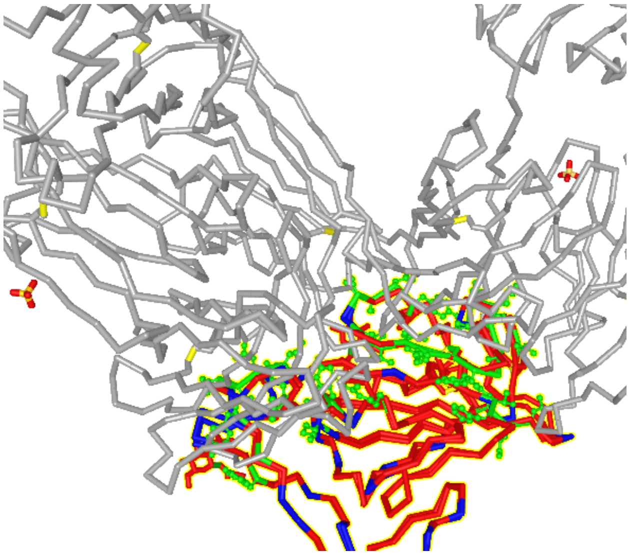

Link 2: PDB 6W41_2DD8: antibodies against SARS-COV-1 vs. SARS-COV-2

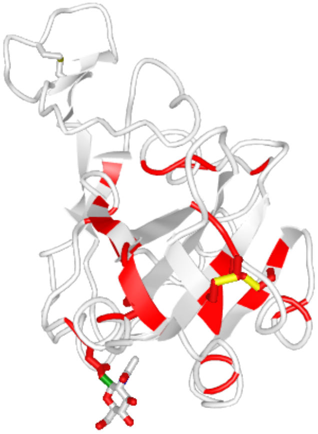

Link 3: PDB 6M0J: mutations may occur across coronaviruses

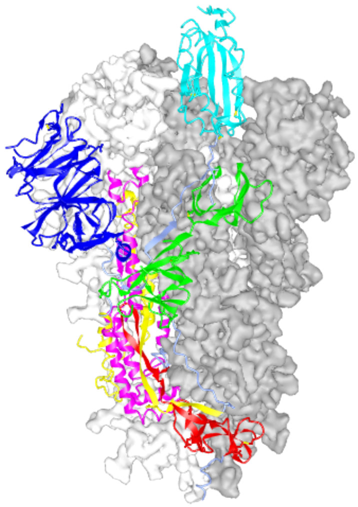

Link 4: PDB 6VSB: domain composition in ribbon representation

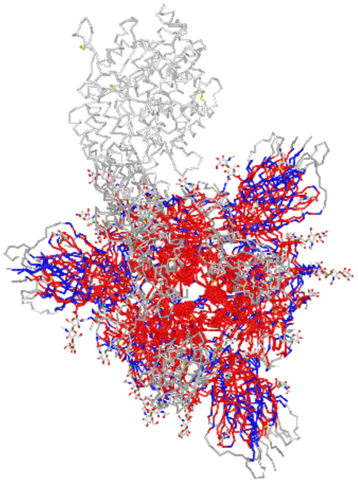

Link 5: PDB 6CS2_6VSB: SARS-COV-1 spike is highly homologous and can be superimposed for the whole trimer (PDB 6CS2)

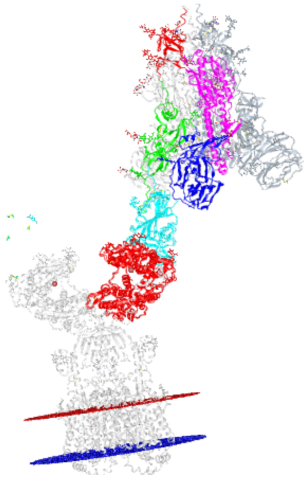

Link 6: PDB 6M17_6CS2: the spike Trimer structure (6CS2) of SARS-COV-1 binding the full length ACE2 peptidase (PDB 6M17)

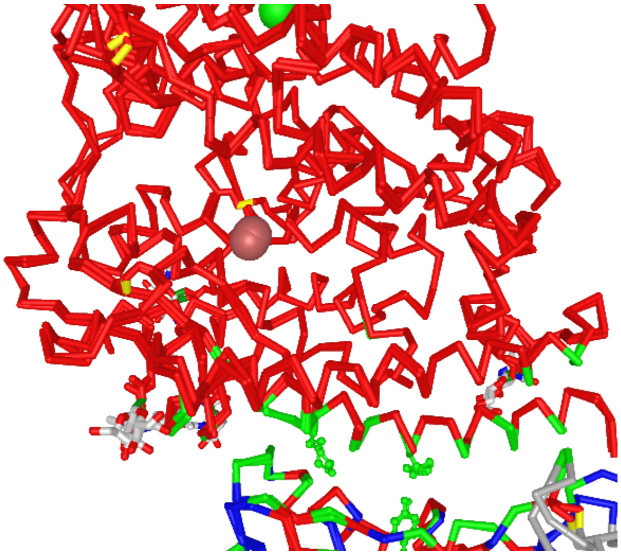

Link 7: PDB 6M0J_2AJF: compare SARS-CoV-1 and SARS-CoV-2 RBD interfaces by superimposing the two complexes (quaternary structures) 6M0J vs. 2AJF

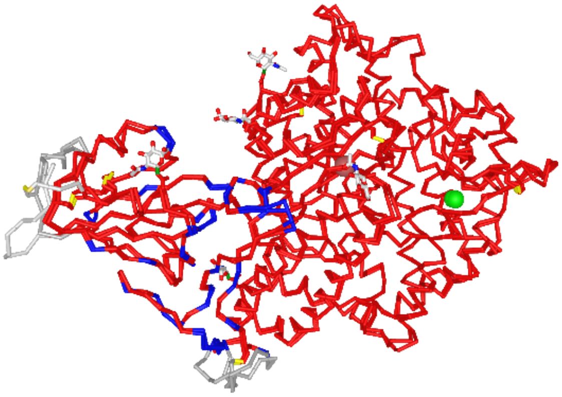

Link 8: PDB 6M0J_3SCJ: compare SARS-CoV-1 and SARS-CoV-2 RBD interfaces by superimposing the two complexes (quaternary structures) 6M0J vs. 3SCI

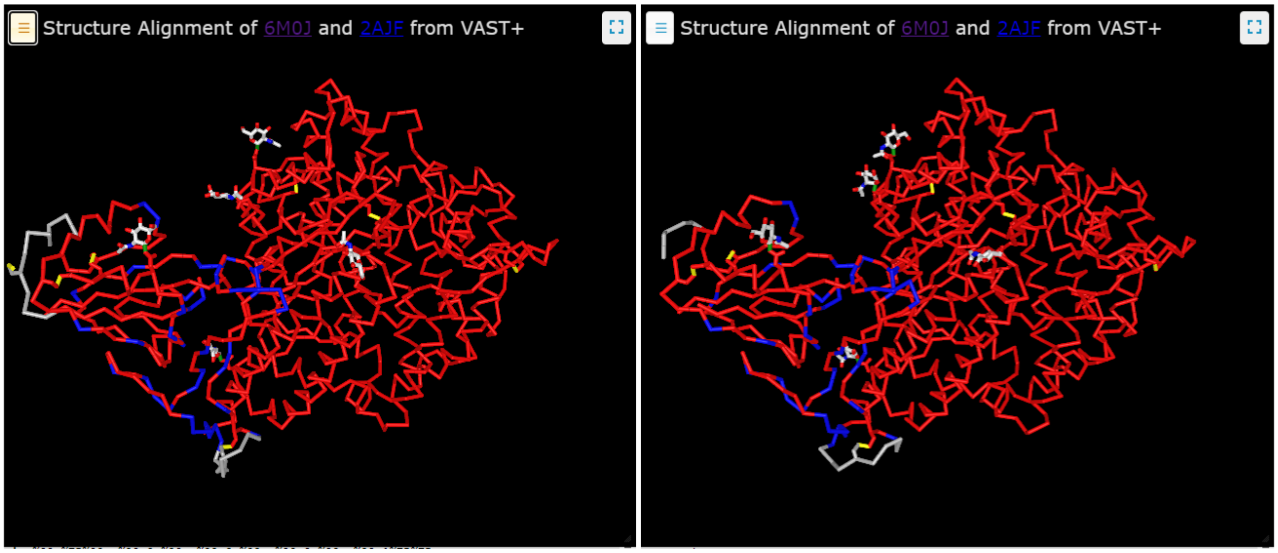

Link 9: PDB 6M0J_2AJF: side by side comparison of SARS-CoV-1 and SARS-CoV-2 RBDs

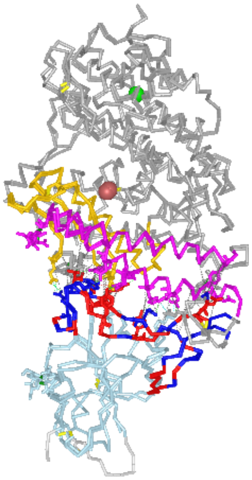
Link 10: PDB 6M0J_2AJF: SARS-CoV-1 - ACE2 vs. the SARS-Cov-2 - ACE2

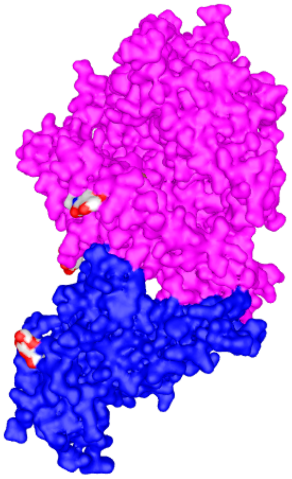
Link 11: PDB 6M0J: the ACE2 - SARS-CoV-2 interface (PDB 6M0J) is formed by two complementary surfaces

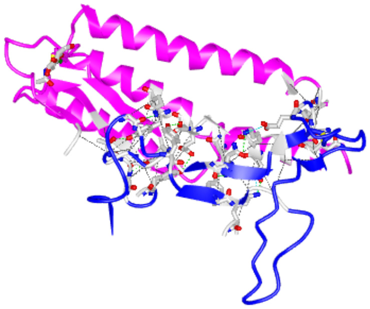

Link 12: PDB 6M0J: ACE2-SARS-CoV-2 interface (PDB 6M0J)

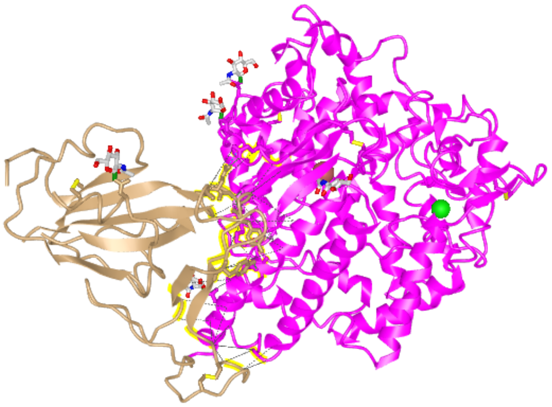

Link 13: PDB 2AJF: ACE2-SARS-CoV-1 interface (PDB 2AJF)

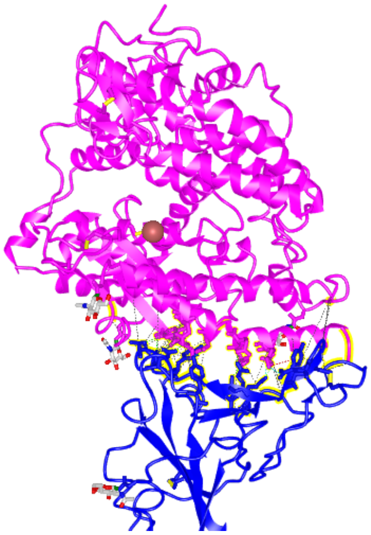

Link 14: PDB 6LZG: the SARS-CoV-2 the interface is composed of 7HBs, 2SBs but also 2 cation-Pi interactions for a total of 40 non bonded interactions

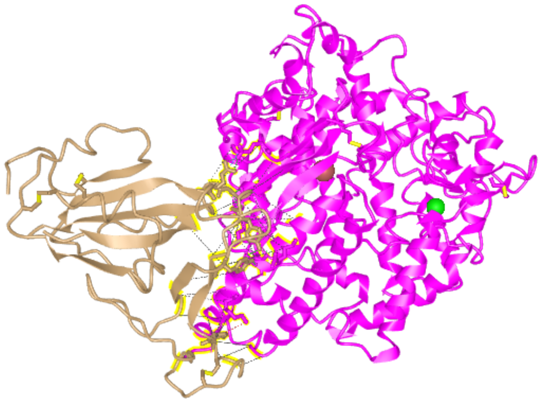

Link 15: PDB 3SCI: the SARS-CoV-1 interface is composed of 5 HBs, 2SBs, 1 CP, 1 PP interactions for a total of 35 nonbond interactions

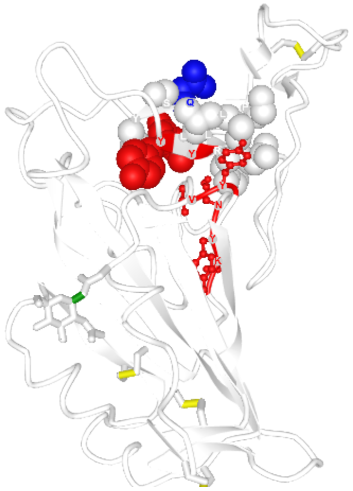

Link 16: PDB 6M0J: highlighting of the YxY conserved motif

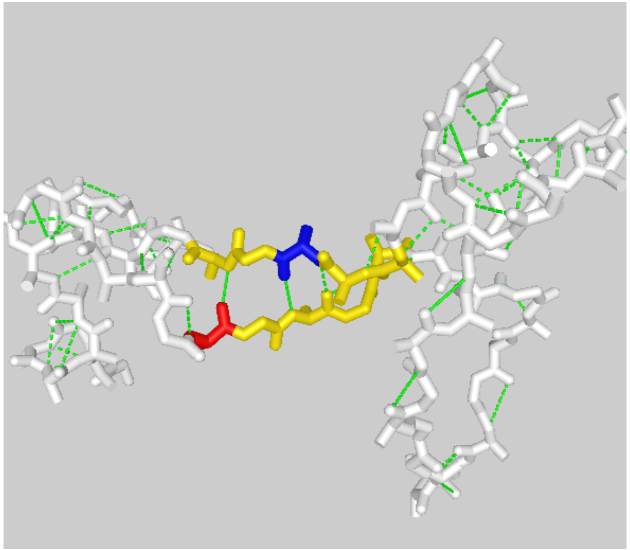

Link 17: PDB 6M0J: RBM supersecondary structure

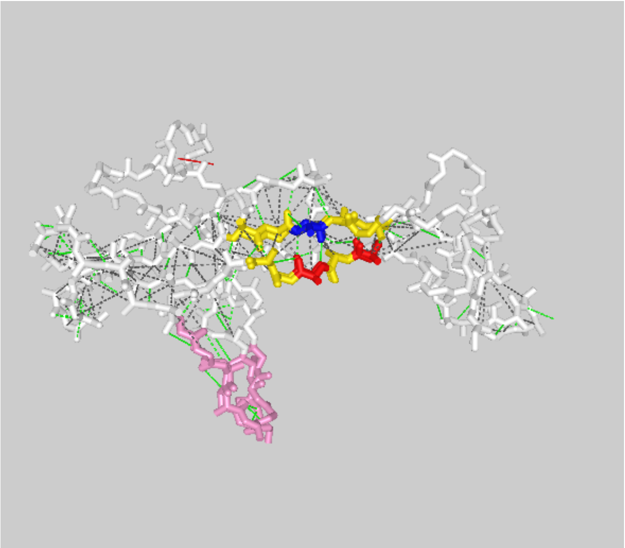

Link 18: PDB 6M0J_4KR0: Comparing MERS to SARS RBM

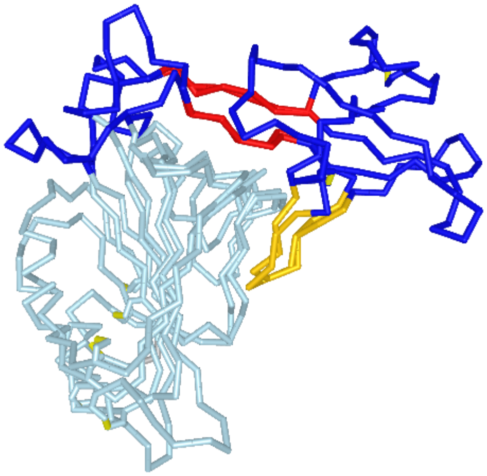

Link 19: PDB 6M0J_4KR0: significant structural variability in the RBM between MERS and SARS viruses

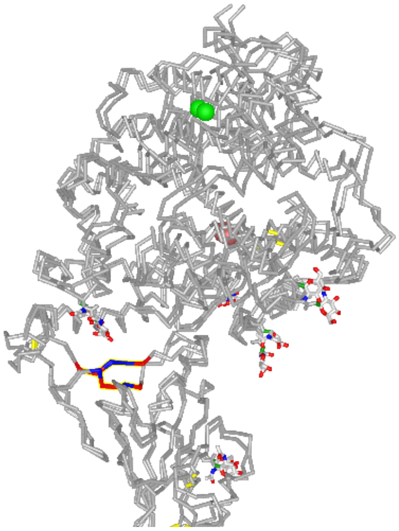

Link 20: PDB 6M0J_2AJF: structurally conserved between SARS-CoV-1 and SARS-CoV-2 with an RMS of 0.304 A

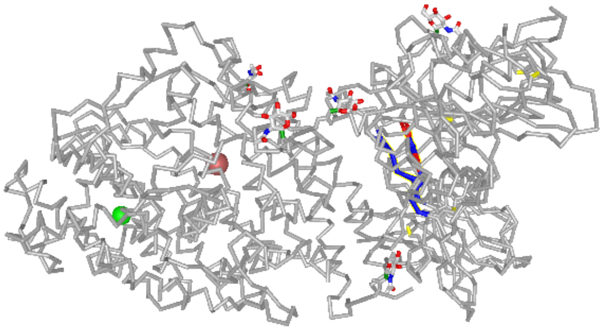

Link 21: PDB 6WAR: structurally conserved vs MERS RBM, where is aligns within 0.64 A RMS

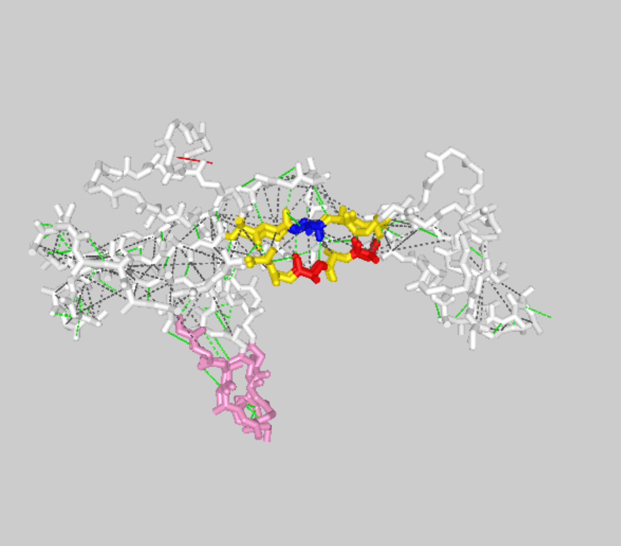

Link 22: PDB 6M0J_4KR0: The 2x5 residue long structural pattern with the YxY sequence motif is conserved in MERS

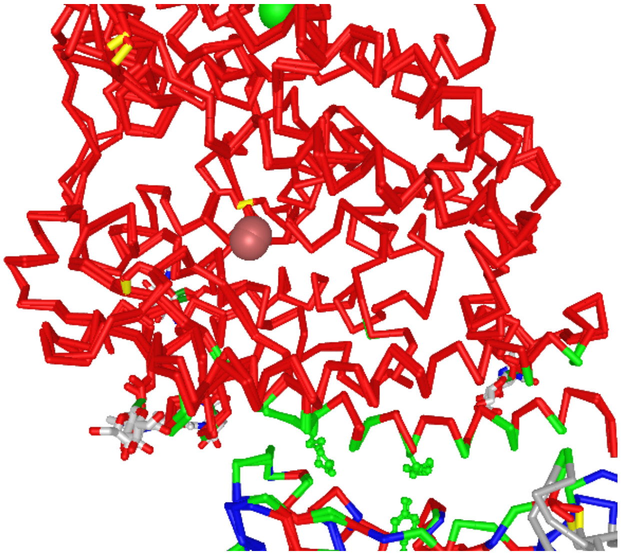

Link 23: PDB 6M0J_2AJF: Structure based sequence alignment of the quaternary complex RBD (A) -ACE2 (B) comparing SARS-CoV-1 and -2

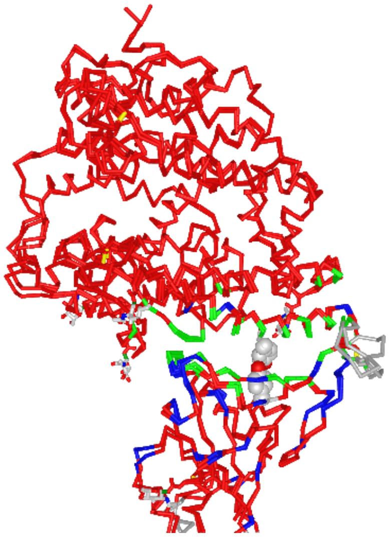

Link 24: PDB 3D0I_6M0J: (6M0J vs 3D0I), again the central RBM-ACE2 interface is conserved overall

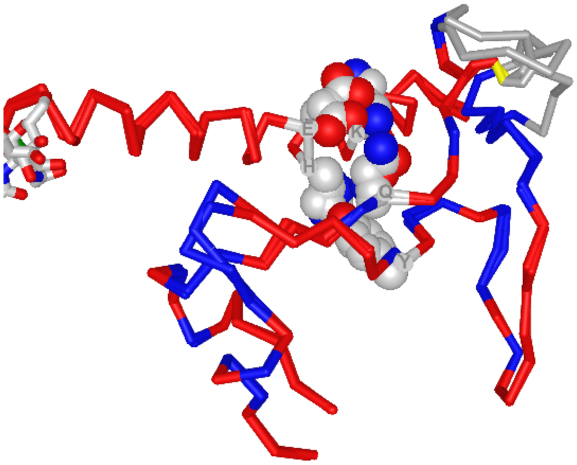

Link 25: PDB 6M0J_2AJF: Comparing SARS-CoV-2 vs. SARS-CoV-1 binding of the RBM-SSS to hACE2

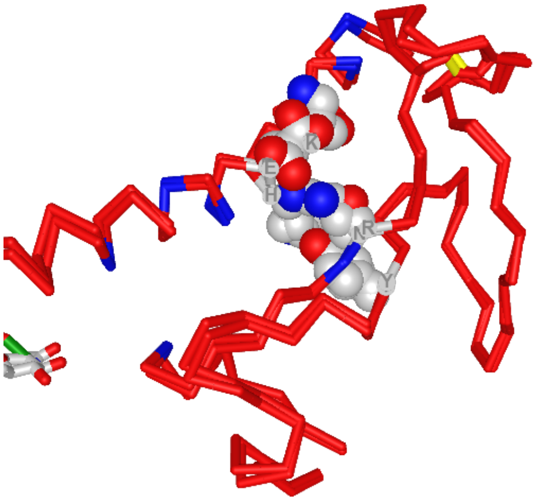

Link 26: PDB 3D0I_2AJF: Comparing Civet-CoV/cACE2 (chimeraN-terminus) vs. SARS-CoV-2/hACE2

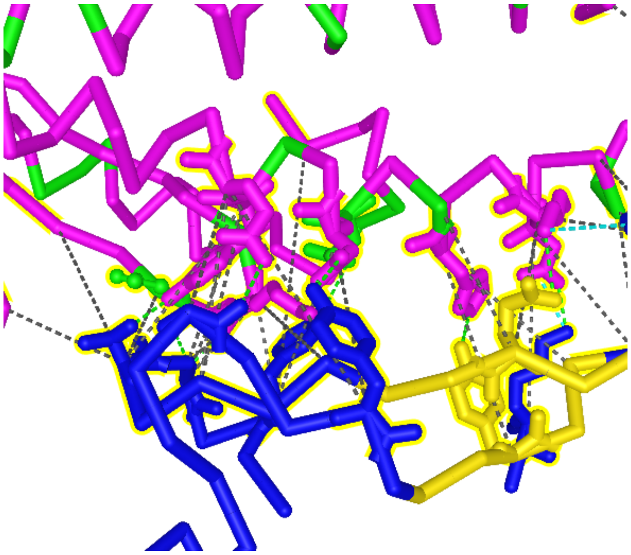

Link 27: PDB 6M0J: ACE2 human polymorphism at the RBD interface

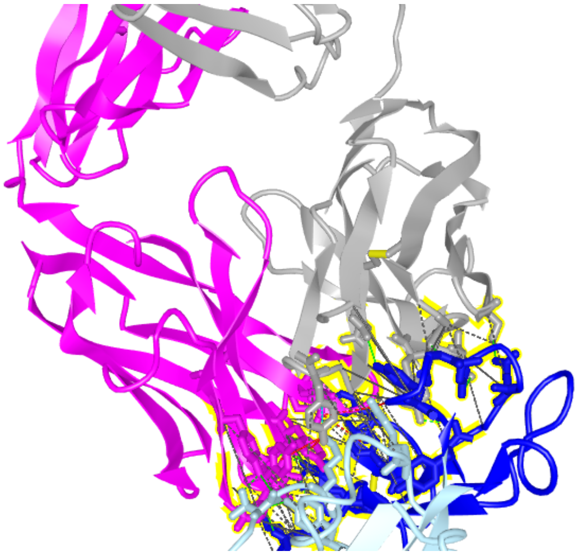

Link 28: PDB 7BZ5: b38 cover the entire RBM of SARS-CoV-2
